# How repeats rearrange chromosomes in deer mice

**DOI:** 10.1101/2024.05.29.596518

**Authors:** Landen Gozashti, Olivia S. Harringmeyer, Hopi E. Hoekstra

## Abstract

Large genomic rearrangements, such as chromosomal inversions, can play a key role in evolution and often underlie karyotype variation, but the mechanisms by which these rearrangements arise remain poorly understood. To study the origins of inversions, we generated chromosome-level *de novo* genome assemblies for four subspecies of deer mice (*Peromyscus maniculatus*) with known inversion polymorphisms. We identified ∼8,000 inversions, including 47 mega-base scale inversions, that together affect ∼30% of the genome. Analysis of inversion breakpoints suggests that while most small (<1 Mb) inversions arise via ectopic recombination between retrotransposons, large (>1 Mb) inversions are primarily associated with segmental duplications (SDs). Large inversion breakpoints frequently occur near centromeres, which may be explained by an accumulation of transposable elements in pericentromeric regions driving SD formation. Additionally, multiple large inversions likely arose from ectopic recombination between near-identical centromeric satellite arrays located megabases apart, a previously uncharacterized mechanism of inversion formation. Together, our results illuminate how repeats give rise to massive shifts in chromosome architecture.

## Introduction

Although genes display remarkable conservation across millions of years of mammalian evolution^1,2^, their organization on chromosomes can be highly variable even on short evolutionary timescales^3^. Karyotypes, the full sets of chromosomes for an organism, provided the first evidence of genomic structural variation. Early cytogenetic studies revealed that chromosome number varies widely across organisms, suggesting that structural variants such as chromosomal fissions and fusions often differentiate species^4–6^. For example, humans have 23 haploid chromosomes, whereas chimpanzees have 24 haploid chromosomes^7^, which is now known to be driven by a fusion of two ancestral primate chromosomes into the human chromosome 2^8^. Other features of karyotypes also highlight structural variation, such as differences in the number of chromosome arms. Chromosome arm number changes when centromeres are re-positioned from the middle to the end of the chromosome or vice versa. Massive genomic rearrangements, such as chromosomal inversions, can drive these changes through moving centromeres between acrocentric and metacentric states^5^. Comparisons of karyotypes suggested that inversions are a prevalent source of structural variation in diverse species^7,9–11^.

Despite longstanding cytogenetic evidence of structural variation, we still know little about how large structural variants arise within the genome^12^. Chromosomal inversions are an important form of structural variant because they suppress recombination and can facilitate adaptive evolution^13^, but have been particularly elusive for two major reasons. First, inversions are challenging to detect with molecular short-read sequencing data because they are balanced polymorphisms that do not change overall gene content^12^. Second, inversion breakpoints frequently occur in highly repetitive genomic regions, which are difficult to assemble^14–16^ (but see ^17,18^). Modern sequencing techniques such as long-read genome sequencing provide new opportunities to study the molecular basis of inversions. This is exemplified by recent long-read based studies in humans, including the telomere-to-telomere assembly of the human genome, which identified an abundance of inversion polymorphisms within humans^19,20^. These studies highlighted the intimate relationship between genomic repeats and chromosomal rearrangements: ectopic recombination between repetitive genomic elements such as retrotransposons or segmental duplications (low copy duplications larger than 1 kb with 90% identity) often drives inversion formation in humans, with mechanisms differing by inversion length^16,19^. However, inversions in humans are modest in size, rarely exceeding 1 megabase (Mb) in length^19^. Thus, how massive inversions (e.g., reported in other animal and plant genomes^21–23)^ arise remains a major open question in genomics.

The deer mouse (*Peromyscus maniculatus*) serves as a model system for studying the molecular basis of inversions. Since the mid-20th century, cytogenetic studies have highlighted widespread karyotypic variation within the species: wild populations of deer mice display metacentric chromosome number ranging from 16 to 40, indicative of many large chromosomal rearrangements, yet a highly conserved total number of chromosomes (2*n* = 48)^24,25^. Recent work has shown that massive inversions underlie much of the karyotypic variation found within deer mice, including re-positioning centromeres from acrocentric to metacentric positions^26,27^. The deer mouse harbors at least 21 mega-base scale inversion polymorphisms, with evidence for multiple inversions spanning over 40 Mb^26^. Furthermore, these inversions are polymorphic *within* the deer mouse species^26^, suggesting their possibly recent evolutionary origin. Thus, the deer mouse provides an opportunity to investigate the formation of massive mammalian inversions.

Here, we generated chromosome-level *de novo* genome assemblies for four subspecies of deer mice, using a combination of PacBio HiFi long-read and Dovetail Omni-C proximity ligation sequencing, to investigate the origins of deer mouse inversions with unprecedented resolution. We identified ∼8,000 inversion polymorphisms between these subspecies, which together affect over 30% of the genome and include 47 large (>1 Mb) inversions. By assembling complex repeats at inversion breakpoints, as well as entire centromeres across multiple haplotypes, we uncovered genetic mechanisms that give rise to intraspecific structural variation in natural mammalian populations.

## Results

### *De novo* genome assemblies for four deer mouse subspecies

Chromosomal inversions are notoriously challenging to characterize due to their size and often highly repetitive breakpoints^16,19^. Thus, to identify inversions and resolve inversion breakpoints in deer mice, we constructed *de novo* chromosome-level genome assemblies for four deer mouse subspecies. To do so, we crossed two pairs of subspecies (*P. m. bairdii* x *P. m. nubiterrae*, *P. m. gambelii* x *P. m. rubidus*) to generate F1 hybrid individuals. We maximized inversion representation by selecting parents based on their genotypes for known inversion polymorphisms^26^. We then sequenced two resultant hybrid individuals (one female, one male) using a combination of PacBio HiFi long-read and Dovetail Omni-C proximity ligation sequencing (Figure 1A). We produced haplotype-resolved genome assemblies for each hybrid, resulting in one assembly representing each of the four subspecies (Figure 1B,C). To ensure correct phasing, we analyzed local ancestry by mapping short-read resequencing data for each subspecies to each respective assembly, and manually adjusted haplotypes when necessary (Supplemental Figure S1). These efforts resulted in highly contiguous chromosome-level assemblies for each subspecies (Supplemental Figure S2; Supplemental Table S1). These *de novo* assemblies show strong improvements to the previous deer mouse reference genome assembly (Supplemental Figure S3), including, for example, a ∼15% increase in repeat representation (Supplemental Table S2; Supplemental Figure S4).

**Figure 1.**
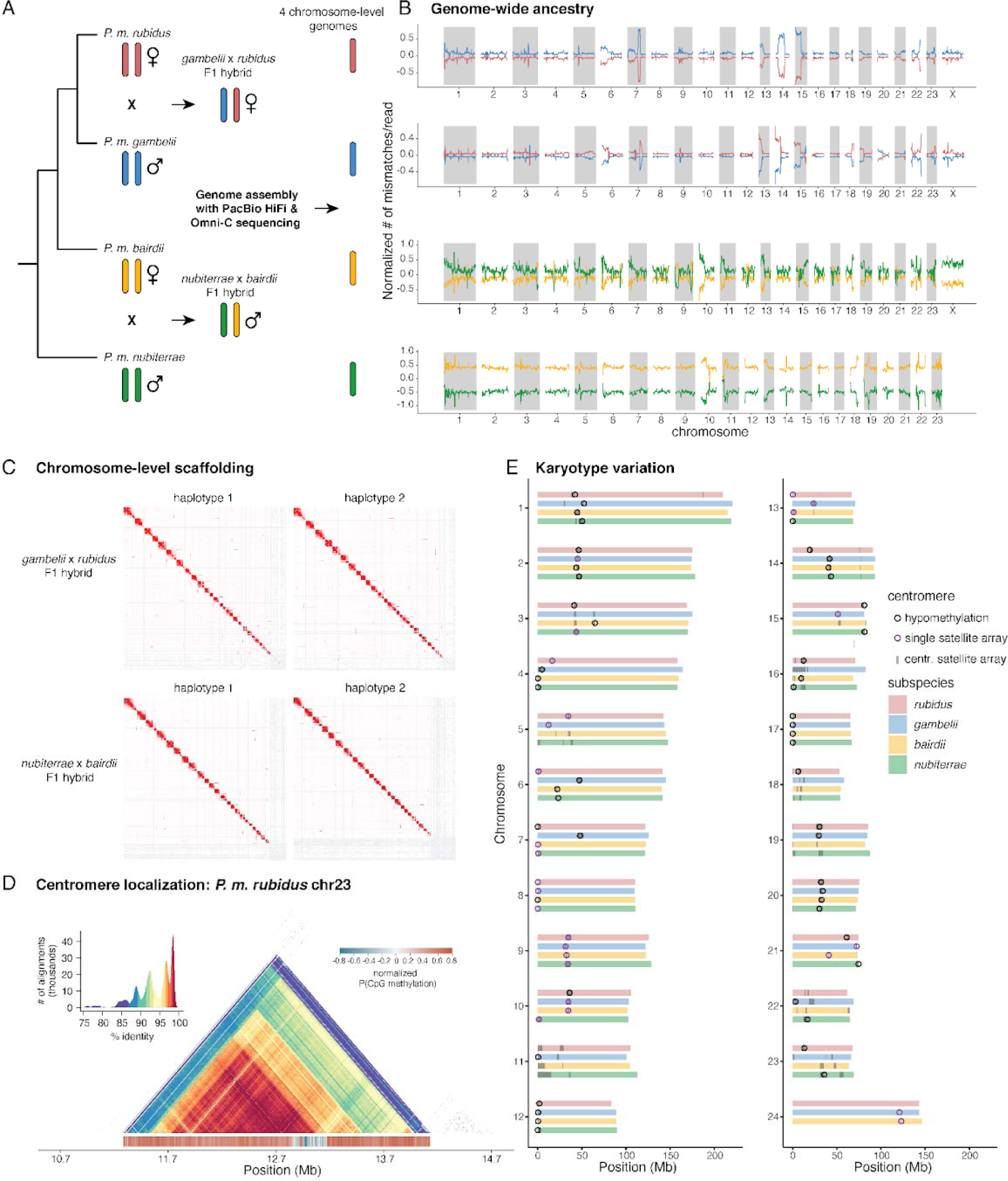
*De novo* genome assemblies and karyotypic variation: (**A**) The genomes of two F1 hybrids were sequenced to generate four chromosome-level genome assemblies. Cladogram of the relatedness between subspecies is shown, with sex indicated for each individual. (**B**) Estimated subspecies ancestry across the genome for each of the four genome assemblies. Colors correspond to subspecies shown in (A). Ancestry is estimated using the normalized number of mismatches per read based on short-read whole-genome sequencing data mapped to each genome assembly (see Methods), with lower values indicating closer ancestry match. Ancestry switches on chr20 and chr21 (*rubidus*) are due to polymorphic inversions within the population-level sequencing data; large peaks correspond to inversions segregating between subspecies. (**C**) Contact maps from Omni-C data for each haploid genome, plotted in Juicer. All genomes have 24 major scaffolds, corresponding to chromosomes, except for the male individual (*nubiterrae* x *bairdii* haplotype 1) with 23 major scaffolds. (**D**) Example of an assembled centromere in the *P. m. rubidus* genome (chromosome 23). Heatmap shows all-by-all percent identity calculated for 5 kb windows. CpG methylation is shown below the heatmap, calculated for 5 kb windows, with the hypomethylated region suggesting the active centromere location. (**E**) Karyotype variation across the four genome assemblies. Centromere positions were estimated based on centromeric satellite sequence locations (gray shaded regions), with hypomethylation patterns localizing centromere positions (black circles); in the absence of clear hypomethylation patterns, centromeres were also localized on chromosomes with only one centromeric satellite array (purple circles).

To evaluate karyotype diversity across the four genomes, we localized centromeres for most chromosomes in each subspecies. We initially inferred centromere locations by mapping previously identified centromeric repeats to each assembly (see Methods)^28^. Although centromeres are often amongst the most difficult regions to assemble because of their high repeat content^29,30^, our *de novo* genome assemblies contained large regions of assembled centromeric satellite sequence on almost every chromosome (Figure 1D,E). We defined potential active centromeres based on the presence of tandem centromeric repeat arrays spanning at least 100 kb^31^. However, many chromosomes harbored multiple, distinct centromeric repeat arrays, varying from 1 to 4 arrays per chromosome with array lengths spanning up to 15 Mb (Figure 1E; Supplemental Figure S5). Since chromosomes with multiple active centromeres are often highly unstable^32–34^, we expected that only one repeat array contained the site of kinetochore assembly on each chromosome and used patterns of CpG methylation to further refine predicted centromere positions. Active centromeres show signatures of DNA hypomethylation at the site of kinetochore assembly^29,30,35^. Thus, to resolve these ambiguities and discern active centromeres, we predicted CpG methylation landscapes using PacBio HiFi kinetics data and searched for regions of reduced methylation within candidate centromeres (Figure 1D; Supplemental Figure S6). Indeed, most chromosomes showed only one distinct region exhibiting centromeric satellite sequence with extensive hypomethylation, thus serving as the predicted centromeres. Based on our localization of centromeres, we found that 13 of 24 chromosomes showed variation in acrocentric and metacentric karyotypes across the four subspecies (Figure 1E). These data suggest that the four genomes show substantial karyotypic variability consistent with observations from early cytogenetic studies in deer mice^24,25^.

### Large chromosomal inversions are common and drive centromere repositioning

Comparison of chromosome-level genome assemblies revealed a wealth of structural variation among deer mouse subspecies. Since *P. maniculatus bairdii* previously served as the representative reference genome for the deer mouse, we called structural variants (>50 bp) with respect to *P. m. bairdii* for the purposes of consistency and comparability^26,27^. Using a combination of *svim-asm*^36^ and *syri*^37^, we identified over 580,000 SVs, including ∼572,000 indels, ∼8,000 inversions and ∼1,000 duplications (Figure 2A,B). Subspecies show similar distributions of lineage-specific and shared SVs, reflecting consistent high-contiguity across assemblies and the utility of each additional genome (Figure 2E). Although fewer in number than indels, inversions affect seven times more of the genome (∼30% for inversions compared to ∼4% for indels; Figure 2C). Consistent with this, inversions represent the largest SVs in deer mice, with 47 inversions covering at least 1 Mb of the genome (Figure 2D; Supplemental Figure S7). This set of inversions includes many previously indicated mega-base scale inversions^26^ and 28 newly described large inversions (Figure 2F).

**Figure 2.**
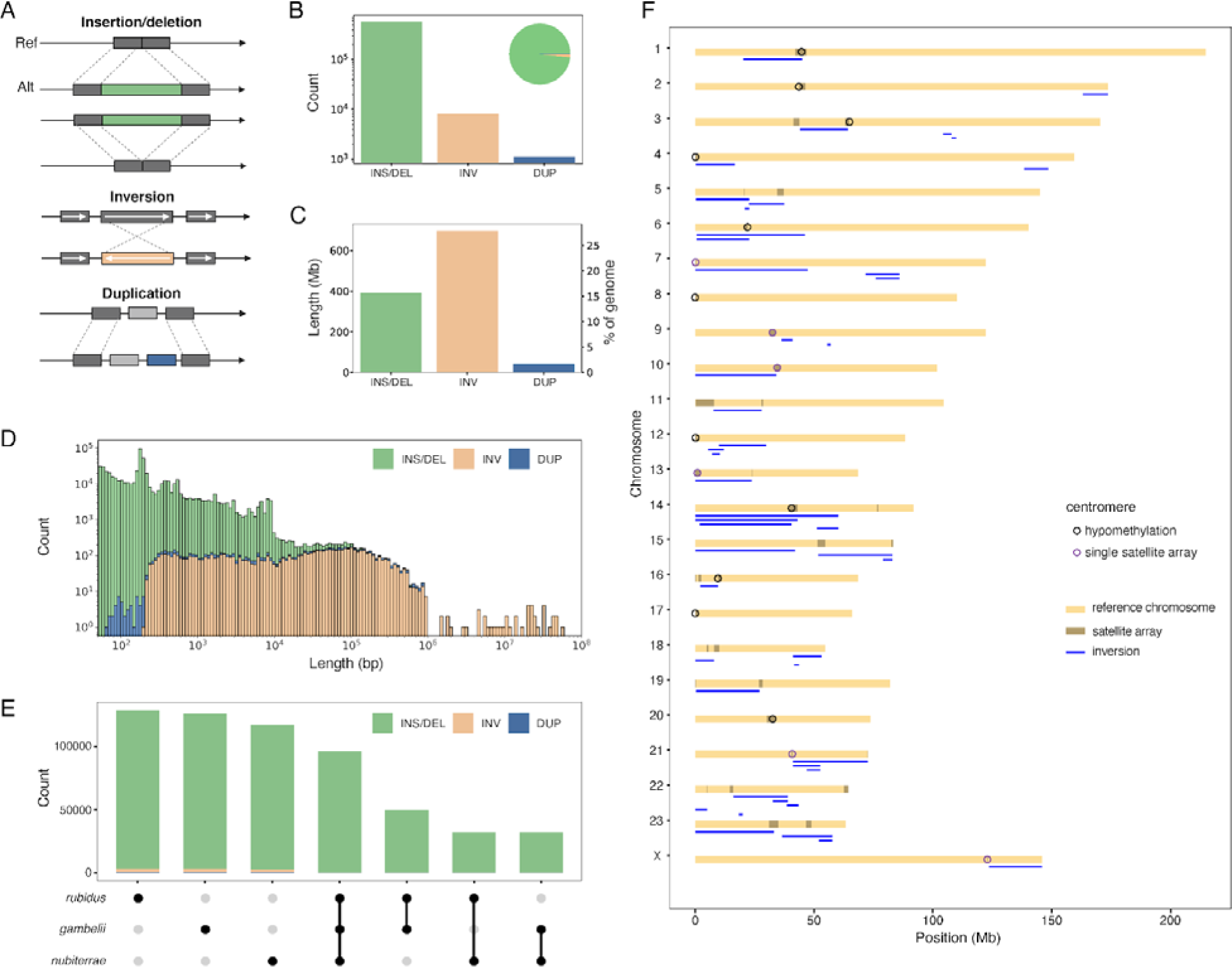
Structural variation includes massive inversions: (**A**) Classes of genomic structural variants (SVs) identified in deer mice. Insertions and deletions are grouped together. (**B**) Number of SVs identified for each class. Pie chart shows proportions of total SVs by class. (**C**) Number of base pairs and percent of the genome affected by each class of SVs. (**D**) Length distributions, shown as stacked bar plots, for each SV class. (**E**) Number of SVs (relative to *P. m. bairdii*) shared by, or unique to, each subspecies. Total number of SVs per subspecies: *rubidus* = 308,426; *gambelii* = 306,130; *nubiterrae* = 278,649. (**F**) Diagram of mega-base scale inversions in the *P. m. bairdii* genome. Inversions are annotated with blue lines. Centromere positions, as predicted by the presence of a single centromeric satellite array or signatures of hypomethylation, are annotated with purple and black open circles respectively.

The megabase-scale inversions are distributed across 21 of 24 deer mouse chromosomes (Figure 2F). Of these large inversions, at least 14 are likely pericentric (contain the centromere) based on our predicted centromere locations. This indicates that these inversions underlie variability in centromere locations across the four subspecies. Furthermore, consistent with previous observations^26^, we found that many (42%) of the mega-base scale inversions exhibit breakpoints within pericentromeric and centromeric regions and ∼40% within 1 Mb of chromosome ends (Figure 2F). These results suggest that inversions play an important role in deer mouse genome evolution and underlie centromere repositioning. Furthermore, the locations of these inversion polymorphisms raise the question of why large inversions tend to have breakpoints near centromeres.

### Large inversions arise repeatedly at the same breakpoints

Our set of mega-base scale inversions included multiple examples of overlapping inversions (including two regions of previously identified overlapping inversions^26^) (Figure 3, Supplemental Figure S8). Some of these regions showed complex nested rearrangements (e.g., chr5, chr14, chr22), which will require future investigation. Nevertheless, our analyses revealed examples of multiple inversions sharing nearly identical breakpoints (Figure 3; Supplemental Figure S8). Using alignments with two outgroup species, we determined the ancestral state for these overlapping inversions and found examples of independently derived inversions sharing the same breakpoint (within 10-kb) (Figure 3A,B). These inversions, such as the 22-Mb and 45-Mb pericentric inversions found on chromosome 6, can contribute to the diverse positioning of centromeres found within the species (Figure 3A). We also identified examples of nested inversions that arose sequentially, with the same breakpoint recurring (Figure 3C). Together, these findings are consistent with the recurrence of inversion breakpoints observed in other species^14,16,19,38,39^; however, while previous studies in great ape lineages showed recurrence of entire inversions, our findings suggest that the same genomic region may be involved in the formation of different inversions.

**Figure 3.**
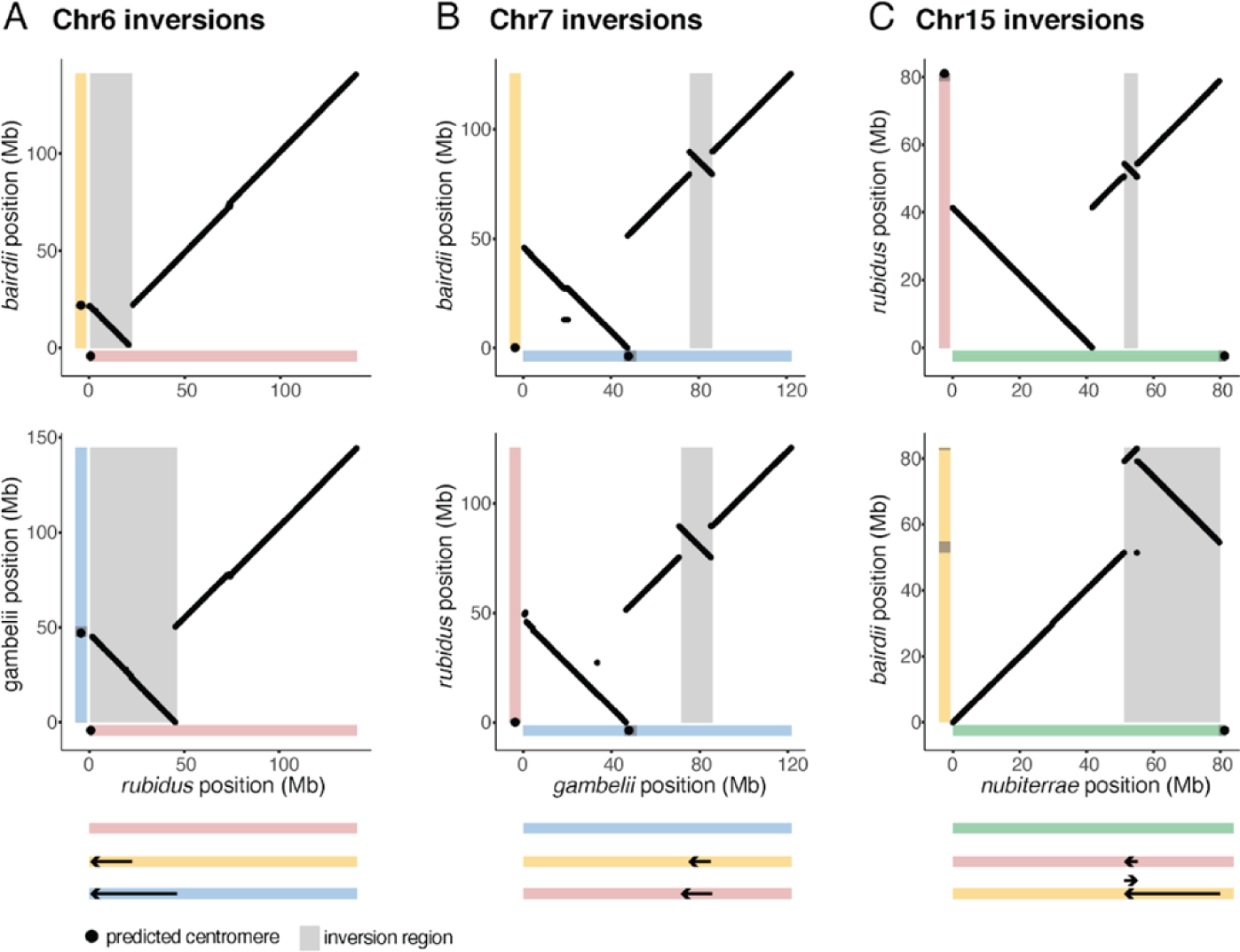
Large inversions share near-identical breakpoints: Examples of inversions with shared inversion breakpoints. Dotplots show alignments (>10 kb) between two subspecies genomes, with ancestral allele on the x-axis and derived allele on the y-axis. Inversions with shared breakpoints are independently derived on chr6 (**A**) and chr7 (**B**). Inversions with shared breakpoints on chr15 are nested (**C**). Diagrams of the inversions with shared breakpoints are shown below dotplots, with chromosomes colored by subspecies and arrows denoting inversions. Additional examples are shown in Supplemental Figure S8.

### Different genomic repeats may drive inversions of different sizes

To investigate the breakpoints of deer mouse inversions, we first characterized genome-wide repeat content. Inversion breakpoints are commonly associated with repeats, since repeats may drive the formation of inversions or arise as an indirect consequence of inversions^15,16,19,40–42^. Thus, to test for associations between inversions and different genomic repeats, we annotated transposable elements (TEs) and segmental duplications (SDs, defined as >1 kb low copy intrachromosomal repeats with ≥90% identity) in the deer mouse genome. We additionally annotated the deer mouse centromere satellite sequence^28^.

We found that inversions showed a strong enrichment for repeats at their breakpoints, but with different patterns based on inversion size. Specifically, we compared repeat occupancy at inversion breakpoints to the rest of the genome and performed permutation tests for enrichment with respect to random expectations. For short (<1 Mb) inversions, we found that both TEs and SDs are enriched at inversion breakpoints; this enrichment includes all three subclasses of active retrotransposons (LINEs, SINEs, and LTRs) (Figure 4A). The breakpoints of short inversions exhibit steep peaks of repeat enrichment that rapidly degenerate, suggesting the presence of localized repetitive regions at these inversion breakpoints (Figure 4B,C).

**Figure 4.**
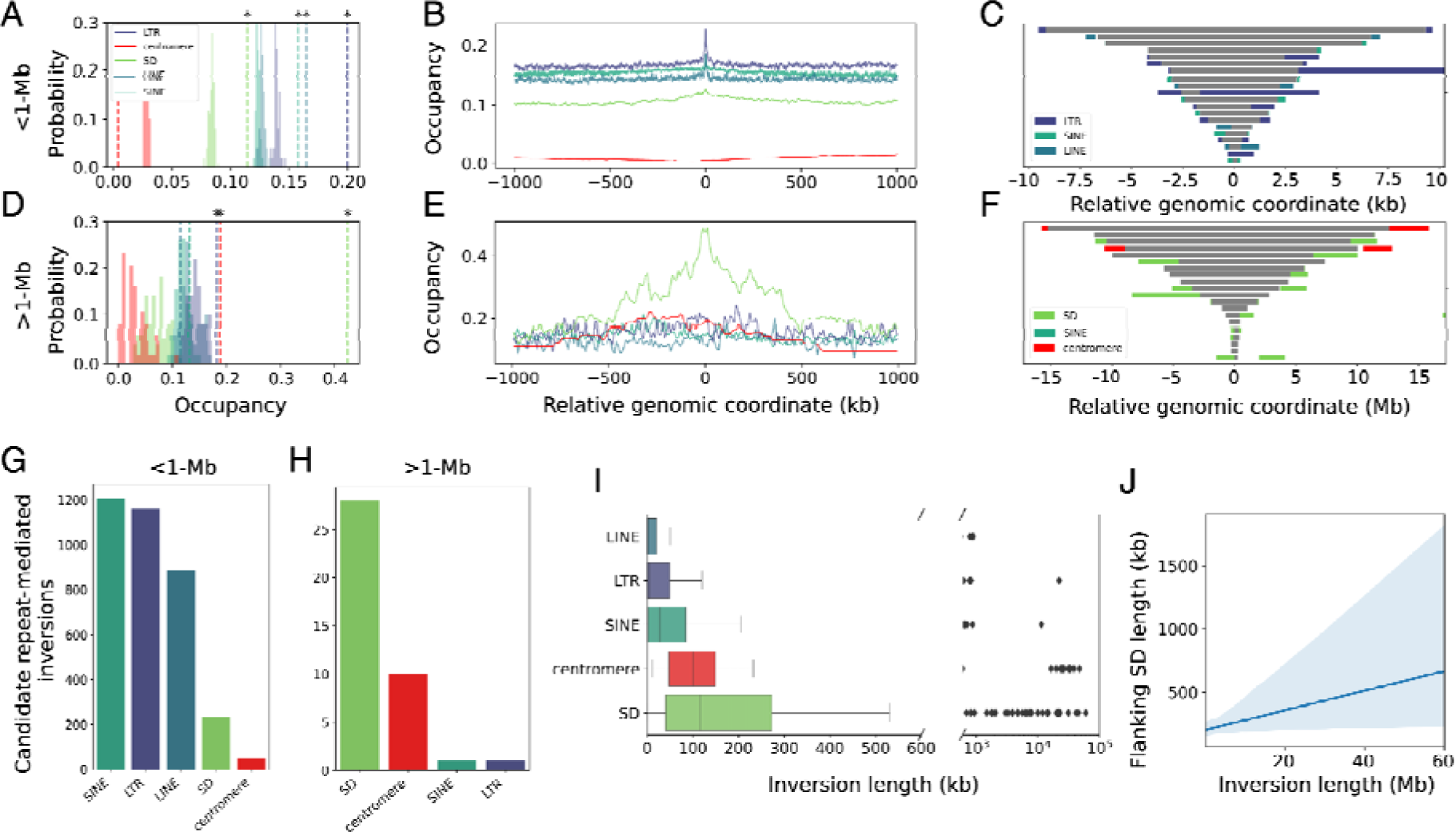
Repeats enriched at inversion breakpoints: (**A**) Expected repeat occupancy at inversion breakpoints from 1,000 random resampled permutations (histograms) compared to observed repeat occupancy (dotted lines) for all inversions <1 Mb. Asterisks denote significant enrichment (P<0.05). (**B**) Average repeat occupancy calculated for 1 kb windows relative to inversion breakpoint position (centered on zero). (**C**) Flanking repeat locations relative to inversion positions for 15 randomly sampled <20 kb inversions (plot inspired by^19^). Inversions are shown as gray bars. (**D**), (**E**), and (**F**) are the same as (A), (B), and (C) respectively but for inversions >1 Mb. (**G**-**H**) Candidate repeat mediated inversion counts for (**G**) inversions <1 Mb and (**H**) inversions >1 Mb. (**I**) Length distributions for inversions associated with each repeat class. (**J**) Linear regression and 95% confidence interval with a “robust” model accounting for outlier effects for average flanking SD length by inversion length^112^ (Kendall’s Tau = 0.216, P<0.00001).

In contrast, TEs are rarely associated with the breakpoints of mega-base scale inversions. We found no evidence for overrepresentation of LINE or SINE retrotransposons at the breakpoints of megabase-scale inversions (Figure 4D). The large (>1 Mb) inversions primarily overlap SDs at their breakpoints: we observed a strong signal of enrichment for SDs, with breakpoints displaying a fourfold increase in SD occupancy with respect to random expectations (Figure 4D). Moreover, SD enrichment extends hundreds of kilobases beyond inversion breakpoints, suggesting that the breakpoints of large inversions occur in complex genomic regions with extended SD content (Figure 4E,F; Supplemental Figure S9). We also observed a strong enrichment of centromeric repeats at the breakpoints of large inversions (Figure 4D-F). Indeed, 47% of large inversions have at least one breakpoint residing *within* centromere satellite arrays (see Figure 2F), demonstrating how inversion breakpoints not only occur near centromeres but also within centromeric satellite arrays.

The enrichment of repeats at inversion breakpoints raised the question of whether these repeats play a role in the formation of inversions. Indeed, inversions commonly arise via nonallelic homologous recombination (NAHR) between repetitive sequences such as TEs or SDs^19,41,43^. To identify candidate NAHR-mediated inversions, we searched inversions for cases in which the same class of repeats was found at both inversion breakpoints^16,19^. For the short (<1 Mb) inversions, we found that ∼47% show homologous TEs at both breakpoints (Figure 4C,G). Among TEs, SINEs and LTR retrotransposons are associated with the most inversions, followed by LINEs (Figure 4G). These results suggest that TEs may play an important role in mediating short inversions.

By contrast, the vast majority of large (>1 Mb) inversions show SDs at both inversion breakpoints, suggesting that SDs, and not TEs, are more likely to contribute to the formation of large inversions (Figure 4F,H). In addition, a subset of large inversions show centromeric satellite sequence flanking both breakpoints, hinting at a possible role of the centromeric satellite sequence in inversion formation (Figure 4F,H). One major challenge in analyzing these large inversions, however, is that the inversion breakpoints often coincide with contig boundaries from the initial *hifi-asm* contig-level assemblies: over 60% of mega-base scale inversions have at least one breakpoint at a contig boundary or chromosome end (Supplemental Figure S7). The coincidence of inversion breakpoints and contig boundaries suggests that these inversion breakpoint regions may not be completely assembled in the genomes (noting that the Omni-C data enabled scaffolding contigs into chromosomes), further underscoring the high repeat content and complexity of these breakpoint regions. Nevertheless, we found that most adjacent contig ends show similar repeat content (Supplemental Figure S10), suggesting that the contig boundaries likely represent the repeat content at inversion breakpoints (if not the full extent of those repeats). Therefore, while we could not identify all causal SDs underlying the formation of massive inversions (but see, Supplemental Figure S9), our results suggest SDs and centromeric satellites likely play a role in inversion formation.

Overall, we found that SD and centromere-associated inversions are significantly longer than TE-associated inversions (Figure 4I; Mann-Whitney U test, P<0.001 for both SD v. TE and centromeric repeats v. TE). This suggests that different repeats may facilitate the formation of inversions of different sizes. Additionally, inversion size and flanking SD size are positively correlated, suggesting that longer SDs may facilitate longer inversions, consistent with observations in humans^19^ (Kendall’s Tau = 0.216; P<0.00001; Figure 4J; Supplemental Figure S11). Together, these results suggest that while TEs primarily give rise to smaller inversions, larger SDs and centromeric satellite arrays are associated with larger inversions in the deer mouse genome.

### Retrotransposons shape SD distributions

Given the association between SDs and megabase-scale inversions, we next investigated the landscape and origins of SDs. We found that SDs are highly enriched near centromeres, which may help explain why inversion breakpoints frequently occur in pericentromeric regions (Figure 5A). To investigate the origins of deer mouse SDs, we searched for TEs at SD breakpoints, since NAHR between TEs can drive segmental duplications^44,45^. The relative age of SDs and TEs can be estimated using divergence between homologous copies or divergence from a respective consensus sequence^46^. Comparison of TE and SD divergence distributions suggests that a large, recent expansion of SDs followed a proliferation of LINE and LTR retrotransposons (Figure 5B). This influx of LINE activity, which was not represented in the previous deer mouse genome assembly^47^, resulted in highly similar LINE copies occupying >1% of the deer mouse genome (Supplemental Table S3). To assess whether TEs could have driven SD expansions, we looked for highly similar TE copies at the breakpoints of SD partners. Moreover, since potentially causal TEs must predate associated SDs, we required that candidate causal TE divergence exceeded associated SD divergence (Figure 5C). We then compared the number of SDs displaying these patterns to expectations from randomization, revealing nearly a three-fold enrichment for LINEs, a less-pronounced but statistically significant enrichment for LTRs (Figure 5D; permutation test, n=100, P<0.01 for both; see Methods), and no enrichment for SINEs (Supplemental Figure S12; P>0.99). Indeed, SDs, including those associated with large inversions, often occupy LINE and LTR-rich regions (Figure 5E). Furthermore, consistent with the rapid decline in NAHR probability with mutation accumulation ^48^, causal TE divergence and SD divergence are directly correlated, suggesting that younger TEs facilitated younger SDs (Figure 5F; Kendall’s Tau = 0.157, P<0.00001). These results suggest that TEs are a fundamental source of SDs in the deer mouse genome.

**Figure 5.**
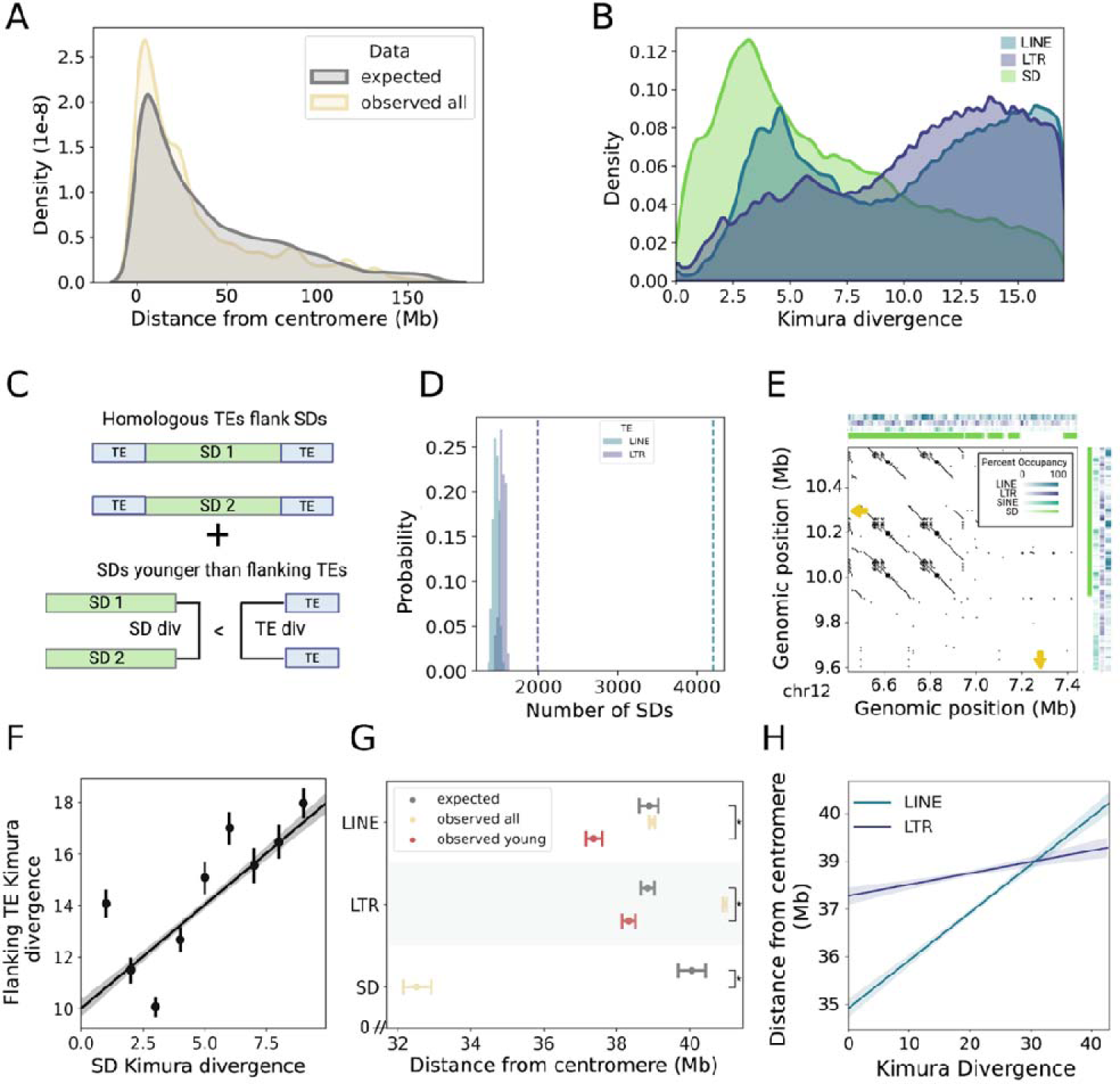
Transposable elements and the evolution of segmental duplications: (**A**) Kernel density estimates for observed distances from the nearest centromere for all SDs compared to expectations from randomization. (**B**) Kernel density estimates for Kimura divergence of LINEs, LTRs, and SDs (including low-copy repeats with <90% identity). (**C**) Expected pattern for TE-mediated SDs. Homologous TEs should flank SD partners and exhibit greater divergence estimates than candidate associated SDs, suggesting that they predated the SDs. (**D**) Number of SDs showing the pattern in (C) compared to expectations from random resampling. Histogram show expected distributions and dotted lines denote observed values. (**E**) Dotplot showing alignments between breakpoint regions for a large inversion on chromosome 12 with flanking inverted segmental duplications. Heatmaps display repeat occupancy calculated across 10 kb windows for each breakpoint. Inversion breakpoints are annotated with yellow arrows. (**F**) Linear regression and 95% confidence interval of flanking TE divergence by SD kimura divergence for all cases in which SDs are flanked by TEs of the same class (Kendall’s tau=0.157, P<0.000001). Data is divided into 9 evenly spaced bins for readability but the regression and confidence interval reflect the original data. (**G**) Observed distributions of distances from th nearest centromeric repeat array compared to expectations from random resampling across different repeat types (asterisk denotes one-sided permutation test, P<0.001). (**H**) Linear regressions and 95% confidence intervals of distance from the nearest centromere repeat by Kimura divergence for different transposable element subclasses (Kendall’s tau=0.0179, 0.0107 (LINEs, LTRs respectively) and P<0.00001 (all)).

Biased accumulation of LINE and LTR retrotransposons in pericentromeric regions may explain SD expansions. While most LINE and LTR retrotransposons are farther from centromeres than expected by chance, young LINEs and LTRs (<10% divergence from the consensus) are significantly closer (Figure 5G; permutation test, n=1000, P<0.001 for both; see Methods). Furthermore, we observe a positive correlation between LINE and LTR age and distance from the nearest centromeric repeat array, suggesting biased accumulation of these elements around centromeres (Figure 5H; Kendall’s Tau=0.0179, 0.0107 and, P<0.00001, <0.00001 for LINEs and LTRs respectively). This bias, and lack thereof for SINEs (Supplemental Figure S12), likely reflects the strong negative fitness effects of LINEs and LTRs, which often fix in AT-rich, gene-poor regions^49,50^. In contrast, SINEs are known to accumulate in GC-rich regions^51^. This biased accumulation of LINE and LTR elements around centromeres may partly explain the enrichment of SDs in pericentromeric regions, which may in turn contribute to the enrichment of inversion breakpoints in these regions.

### Centromeric repeat expansions contribute to large inversions

While pericentric SDs likely drive large inversions in deer mice (Figure 6A), we also found that many massive inversions are flanked by centromeric repeats (Figure 4F,H), suggesting that centromeric repeats themselves may be involved in inversion formation. Two hypotheses could explain the association of centromeric satellite arrays with inversion breakpoints. On the one hand, inversions could facilitate the evolution of new centromeric repeat arrays if one breakpoint occurs within a centromere, moving some but not all of the centromeric satellite array to another chromosomal location (Figure 6B; “centromere-splitting” hypothesis)^52,53^. On the other hand, centromeric satellite arrays could arise in new chromosomal locations via alternative mechanisms and could serve as a potent substrate for large inversions thereafter (Figure 6C; “centromere-mediated” hypothesis)^32,52^. Several centromeric repeat arrays display presence/absence variation among deer mouse subspecies (Supplemental Figure S5). The presence of inverted centromeric repeats at ancestral inversion breakpoints would support the “centromere-mediated” hypothesis that ectopic recombination between centromeric repeats may contribute to inversion formation in deer mice.

**Figure 6.**
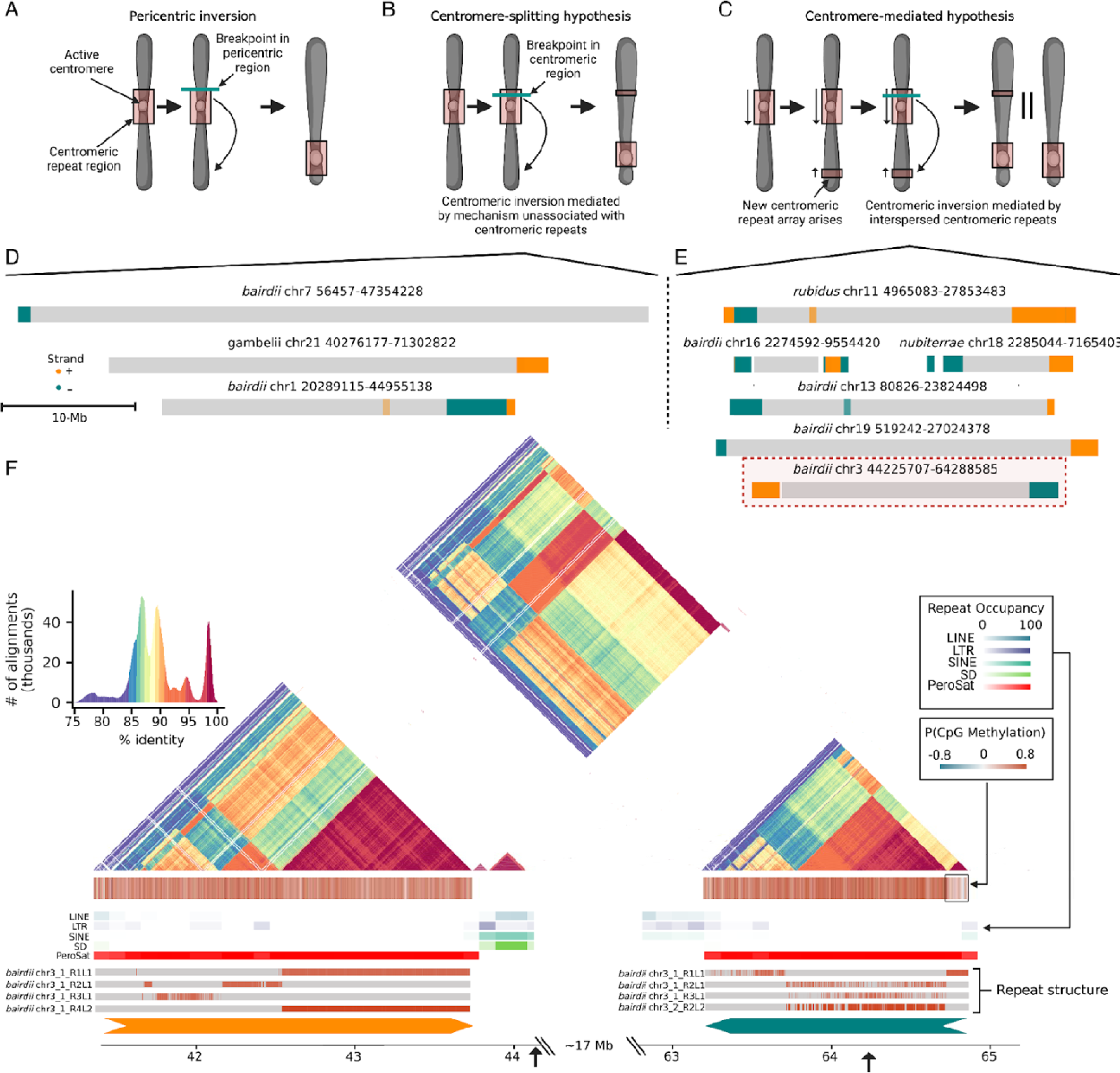
Centromeric satellite arrays mediate massive inversions: (**A**-**C**) Three non-mutually exclusive hypotheses could explain centromere toggling in deer mice. (**A**) A pericentric inversion with breakpoints outside of the centromeric satellite array captures and moves the centromere. (**B**) An inversion with one breakpoint occurring within a centromeric satellite array results in multiple centromeric satellite arrays (centromere-splitting hypothesis). (**C**) An inversion with both breakpoints occurring within centromeric satellite arrays may arise via ectopic recombination between centromeric satellite arrays (if satellite repeats are inverted) (centromere-mediated hypothesis). Depending on the site of ectopic recombination, this mechanism could (but not necessarily) reduce the number of centromeric repeat arrays on a chromosome. (**D**-**E**) Ancestral haplotypes for inversions that display centromeric satellite array (>1 kb) within 1 Mb of both breakpoints, and patterns consistent with either (**D**) the centromere-splitting hypothesis or (**E**) the centromere-mediated hypothesis. Inversions are shown as gray rectangles and centromeric satellite positions are annotated with orange or teal depending on their orientation. Breakpoints for an inversion on chromosome 3 displaying patterns consistent with the centromere-mediated hypothesis (C) (highlighted with a red box) are shown in expanded detail in (**F**). (**F)** In order from top to bottom, heatmap displaying all-by-all percent identity calculated for 5 kb windows^108^; normalized adjusted probability of CpG methylation calculated for 5 kb windows (see Methods); heatmaps displaying repeat density for different repeat classes; distributions for the most common higher order repeat (HOR) structures in chromosome 3 centromeres^54^. HOR repeat naming conventions are structured as follows: RxLy where x is the ranking of repeat coverage (with 1 covering the most bases in the centromere) and y is the HOR level (with 1 denoting monomers and no HOR structure, 2 denoting dimers and so on)^54^. Arrows parallel to genomic coordinates denote centromeric repeat orientation. Black arrows denote prospective inversion breakpoints.

To address these hypotheses, we first identified the ancestral haplotype for each large (>1 Mb) inversion based on alignments with two outgroups (*Peromyscus leucopus* and *Onychomys torridus*). We then analyzed both the ancestral and derived haplotypes for centromeric sequence relative to inversion breakpoints. We found that 9 inversions had centromeric repeats at both breakpoints in at least one of the ancestral or derived haplotypes (excluding recurrent inversions and complex nested inversion regions [i.e., chr5, chr14, chr22]). Of these 9 inversions, 3 were flanked by centromeric repeats in the derived haplotype only, consistent with the “centromere-splitting” hypothesis (Figure 6D). Surprisingly, 6 inversions had centromeric repeats at both breakpoints in the *ancestral* haplotype (Figure 6E). Furthermore, we analyzed the direction of repeats at the breakpoint regions of these inversions in the ancestral haplotypes, and found that all 6 inversions showed inverted, flanking centromeric repeats in the ancestral inversion haplotype (Figure 6E). These analyses provide evidence supporting the “centromere-mediated” hypothesis, highlighting the prospective origins of several inversions as a result of ectopic recombination between centromeric satellite arrays.

For example, an inversion on chromosome 3 has both breakpoints occurring at inverted centromeric repeats with nearly 100% identity spanning >100 kb in the ancestral haplotype (*P. m. bairdii*) (Figure 6F). We also annotated higher order repeat (HOR) structure and monomer subfamily distributions for centromeric repeats at inversion breakpoints^54^. Unlike most mammals^31,55^, deer mouse centromeres appear to lack higher order repeat structures and are primarily composed of highly similar monomeric repeats (Figure 6F; Supplemental Figure S6; Supplemental Table S4). The lack of HOR structure and high sequence similarity in these regions may increase the chances of ectopic recombination. Together, these results suggest that centromeric repeats may contribute to inversion formation.

## Discussion

Chromosomal inversions can have profound effects on the evolution of species. Inversions dramatically affect recombination: when an inversion is heterozygous, recombination is suppressed across the inversion region due to alignment issues between haplotypes^13,56^. Through suppressing recombination, an inversion may link adaptive alleles together into a single, co-inherited haplotype (or supergene), a powerful mechanism to facilitate adaptation^13,15,56^. Indeed, our previous work showed how multiple inversions in deer mice contribute to local adaptation^26,27^: for example, a 41 Mb inversion acts as a supergene linking two forest-adaptive traits together^27^. Inversions can also lead to speciation if they lock together alleles associated with reproductive isolation^57,58^. In addition, inversion breakpoints themselves can cause phenotypic changes through disrupting genes or gene expression^15,56^. Given their importance, we aimed to understand how inversions arise within species. Specifically, we used long-read and proximity ligation sequencing approaches to identify and characterize a diversity of inversions in four deer mouse genomes. Through analyses of inversion breakpoint regions, we revealed the origins of inversions of differing size and illuminated the complex genomic processes shaping their evolution.

We first found that smaller (<1 Mb) inversions likely arise frequently from retrotransposon-associated mechanisms, consistent with observations in diverse species^19,40,43,59–62^. Retrotransposons can generate inversions via ectopic recombination, where two inverted homologous TEs align within a chromosome and NAHR between them results in an inversion^15,59,63^. Highly homologous TEs are more likely to engage in NAHR, which suggests that young TEs may be important for inversion formation^64,65^. Indeed, the deer mouse genome experienced a large invasion of LTR elements^47^, as well as recent LINE/SINE activity evidenced by the *de novo* genomes. These recent TE expansions may play an important role in the formation of small inversions via NAHR across the genome, although TE insertions into breakpoint regions following inversion formation likely also contribute to observed patterns^42^.

In contrast to smaller inversions, large (>1 Mb) inversions are rarely directly associated with TEs. Instead, the majority of mega-base scale inversions we identified are associated with SDs. Breakpoints for these inversions are often highly complex, with SDs extending for tens or hundreds of kilobases, underscoring the importance of long-read sequencing for assembling inversion breakpoint regions^66^. In humans, SDs have been well-documented at inversion breakpoints, with longer SDs associated with longer inversions^16,19^. Consistent with these studies, we found that SD length is strongly correlated with inversion length, suggesting that larger SDs may facilitate larger inversions. This observation may be due to a higher probability of long repeats finding each other over large genomic distances, and/or the necessity for longer stretches of sequence similarity to stabilize the recombination complex between distant chromosomal regions^67,68^. We also found that deer mouse SDs are closer to centromeres than expected by chance, consistent with findings in humans^68,69^. This enrichment is likely driven by selection against SDs in gene-rich regions. However, there may also be a higher probability of SD formation in pericentromeric regions in deer mice. Unlike human SDs, which are mediated by Alu SINE elements that accumulate in GC-rich regions^44,49,69^, we found that deer mouse SDs often arise from LINE and LTR elements, which show recent accumulation in AT-rich, gene poor regions^49,50^. Together, these features of deer mouse SDs may help explain the enrichment of mega-base scale inversion breakpoints near centromeres, noting that selection on inversion breakpoints themselves may also affect the distribution of breakpoints^57^.

While a majority of mega-base scale inversions are associated with SDs, we also found nine massive inversions flanked by centromeric satellite arrays. An interesting feature of the deer mouse genome is that it harbors multiple large (>100 kb) and highly similar (>90% identity) centromeric satellite arrays within single chromosomes, a phenomenon previously observed within the species^28^ and confirmed by our high-quality reference genomes. Multiple large centromeric satellite arrays have rarely been reported in other species^29,70–72^. However, because such centromeric satellite arrays are typically the most difficult regions of the genome to assemble^29,30^, it remains unclear whether this feature is rare or common in the genomes of other species. We initially hypothesized that these multiple deer mouse centromeric satellite arrays arose as a consequence of large inversions re-positioning the centromeric satellites^28,52,73^. However, we found that a majority of inversions flanked by centromeric satellite arrays were likely mediated by ectopic recombination between distant inverted satellite arrays (or recombination between arrays of misaligned homologous chromosomes^68,74^). This is, to our knowledge, amongst the first evidence of centromeric satellite arrays driving chromosomal inversions.

Centromeric satellites have the potential to serve as exceptionally potent drivers of genomic structural rearrangements for several reasons. First, satellite arrays generally evolve through replication slippage, which can lead to rapid repeat expansions while repeat similarity and structure is preserved^29,75–80^. Second, centromeric satellite arrays can be exceptionally long, spanning hundreds of kilobases and even megabases of sequence^31^. Such long arrays of highly similar sequences can thus serve as ideal genomic regions for ectopic recombination. Indeed, deer mouse centromeric satellite arrays located megabases apart are primarily composed of highly similar monomeric repeats with little higher-order repeat (HOR) structure^28^. This feature may help explain the potential of these satellite arrays to serve as substrates for ectopic recombination and subsequent hotspots for inversions. Why deer mouse centromeres show remarkable conservation and minimal HOR structure remains an intriguing question, since these features in many other species exhibit rapid turnover as a result of molecular drive^32,55,81–84^. Although centromere-splitting inversions may account for new centromeric satellite positions on a subset of chromosomes, understanding how multiple centromeric satellite arrays arose within other deer mouse chromosomes will be an important direction for future studies. Several mechanisms for the origins of new centromeric satellite positions have been proposed, including replication of extrachromosomal circles of tandem repeats by a rolling-circle mechanism and reinsertion^85,86^ and transposable element-mediated mobilization^87^. Finally, gene-poor centromeric satellite arrays may serve as safe havens for inversion breakpoints^28,78^, suggesting that selection may also favor inversions occurring within these regions.

Centromeric regions are commonly involved in translocations or fusions due to high interchromosomal conservation at centromeres^32,88^. Despite the high conservation of deer mouse centromeric satellite arrays, interchromosomal rearrangements appear rare within deer mice. Deer mice have a strongly conserved chromosome number (2*n* = 48) both within the species and across the *Peromyscus* genus^89,90^. This is in contrast to many other rodent lineages in which chromosomal fissions and fusions are common^4,91–93^, thus raising the question of why long and highly homologous centromeric satellite arrays seem to facilitate inversions but not translocations and fusions in deer mice. Our findings thus motivate future studies of the role of centromeric satellite arrays, across a diversity of species, in the evolution of inversions and structural variation.

Together, our study demonstrates the intricate relationship between repeats and rearrangements. Although chromosomal rearrangements including inversions have been visible in karyotypes for many decades, their breakpoints have remained obscure. Repeat-rich genomic regions are the most difficult parts of the genome to characterize, and we only recently are able to access these genomic regions ^31^. In uncovering these hidden repetitive parts of the genome, we reveal how small repeats and their expansions can lead to massive chromosomal changes. Moreover, by better understanding the formation of inversions, we gained insight into the mutational processes that influence genome architecture, recombination and, in some cases, organismal fitness.

## Methods

### Genome assembly

To investigate intraspecific structural variation, we first created *de novo* chromosome-level genome assemblies for four deer mouse (*P. maniculatus*) subspecies. Our previous work showed that inversion polymorphisms were common within deer mice and frequently segregated among subspecies^26^. To maximize the number of large (>1 Mb) inversion polymorphisms represented in the genome assemblies, we selected for sequencing four available subspecies with known differences in inversion haplotypes: *P. m. rubidus, P. m. gambelii, P. m. bairdii,* and *P. m. nubiterrae*^26^. We then genotyped 3 mice per subspecies for 9 previously identified large inversions that were polymorphic within subspecies^26^ to further maximize inversion representation. To do so, we designed custom Taqman genotyping assays for diagnostic SNPs for each inversion (Supplemental Table S5). Genotyping reactions were performed with 1-10 ng of genomic DNA from mouse ear clips, using the following cycling parameters: 95°C for 10 minutes followed by 40 cycles of 95°C for 15s, 60°C for 1 minute. We selected mice based on inversion genotypes and sex. We then interbred the interfertile pairs of subspecies (*P. m. rubidus* female x *P. m. gambelii* male, *P. m. bairdii* female x *P. m. nubiterrae* male) to create two first-generation (F1) hybrids. Using F1 hybrids allowed us to improve genome phasing and reduce the cost to create four chromosome-level genome assemblies.

We assembled genomes using a combination of PacBio long-read sequencing and Dovetail Omni-C sequencing, performed by Dovetail Genomics. For PacBio sequencing, we extracted high-molecular weight DNA from fresh whole blood samples (immediately flash frozen) of the two F1 hybrid mice using a Blood and Cell Culture Mini Kit (Qiagen, GmbH) following the manufacturer’s protocol. We quantified DNA samples using Qubit 2.0 Fluorometer (Life Technologies, Carlsbad, CA, USA) and constructed PacBio SMRTbell library (∼20 kb) for PacBio Sequel using SMRTbell Express Template Prep Kit 2.0 (PacBio, Menlo Park, CA, USA) following the manufacturer recommended protocol. We used the Sequel II Binding Kit 2.0 (PacBio) to bind the library to polymerase and loaded libraries onto PacBio Sequel II. We performed sequencing on PacBio SMRT cells, with 5 HiFi SMRT cells per mouse, yielding ∼410-490 Gb/cell with mean subread lengths of ∼12-13 kb. For Omni-C sequencing^94^, we extracted DNA from flash frozen liver, brain and muscle tissues for each sample using Dovetail Genomics’ mammalian tissue sample preparation protocol (see https://dovetailgenomics.com/wp-content/uploads/2021/09/Omni-C-Protocol_Mammals_v1.4.pdf). To prepare each Dovetail Omni-C library, we fixed chromatin in place with formaldehyde in the nucleus. We digested fixed chromatin withLDNaseLI, repaired and ligated extracted chromatin ends to a biotinylated bridge adapter, and then performed proximity ligation of adapter containing ends. After proximity ligation, we reversed crosslinks and purified the DNA. We then treated purified DNA to remove biotin that was not internal to ligated fragments. We generated sequencing libraries using NEBNext Ultra enzymes and Illumina-compatible adapters, and isolated biotin-containing fragments using streptavidin beads before PCR enrichment of each library.LWe sequenced the libraries using Illumina Novaseq platformL to produce ∼ 30x sequence coverage.

Draft genomes were initially assembled using *hifi-asm integrated Hi-C* by Dovetail Genomics with default parameters. This approach creates contig-level genome assemblies using PacBio HiFi data, while integrating Omni-C data to phase contigs into haplotypes^95,96^. The *hifi-asm* assemblies yielded contig N50s ranging from 27.3 to 31.8-Mb (Supplemental Figure S2). We then scaffolded contigs into chromosomes using the Omni-C data. First, we sorted the Omni-C data by haplotype to facilitate subspecies-specific scaffolding of contigs; this step was particularly important for studying large chromosomal inversions that differ in orientation between subspecies. To sort the Omni-C data, we used bwa mem^97^ to map the Omni-C data for each F1 hybrid to a combined genome of its two haplotype-resolved genomes. We mapped paired end reads separately to ensure independent mapping of each read. We then used samtools (version 1.10)^98^ to sort and index bam files and remove any unmapped reads. Next, we selected read pairs for which at least one of the reads uniquely mapped to one haplotype-resolved genome, and the paired read did not uniquely map to the other haplotype-resolved genome, based on contig labels of assigned haplotype. We defined unique mapping as reads with 150 bp aligned to only one of the two haplotype-resolved genomes, with no mismatches; we identified such reads using grep from the bam files. We then selected these uniquely mapping reads using seqtk subseq (version 1.2; https://github.com/lh3/seqtk) from the original Omni-C fastq files. As a result, we obtained ∼96-113M reads per haplotype-resolved genome, corresponding to ∼10-12X coverage. Using these genome-specific Omni-C reads, the haplotype-resolved draft genomes were scaffolded with HiRise by Dovetail Genomics^99^. Finally, Omni-C maps were manually inspected with juicer^100^, and scaffolds were manually joined into chromosomes by Dovetail Genomics (Supplemental Figure S2).

We next performed quality control on the scaffolded genome assemblies. First, we aligned the four assemblies to the Pman2.1.3 genome using minimap2 (version 2.9) with -cx asm5^101^. We found that each of the 24 major scaffolds (or 23 in the case of the male) per genome uniquely aligned to a single chromosome from the Pman2.1.3 genome (e.g., Supplemental Figure S3); we then re-named scaffolds to chromosome number. To ensure that the genomes were correctly phased by subspecies, we took advantage of the divergence between subspecies to analyze ancestry across each scaffold. Specifically, we used whole-genome short-read resequencing data from 15 samples per subspecies (NCBI PRJNA688305, PRJNA838595, PRJNA862503) to select ancestry-informative SNPs. Using a vcf of variants that were previously called using GATK (see ^26^ for details), we identified SNPs with F_ST_ > 0.3 (*rubidus* x *gambelii*) and > 0.5 (*bairdii* x *nubiterrae*) between the two subspecies and recorded allele frequencies by subspecies at each of these SNPs. We then selected these sets of ancestry-informative SNPs from the genome assemblies. To do so, we converted the minimap2 alignments to bam files using samtools sort, and extracted the allele at each of the ancestry-informative SNPs from the bam file using samtools mpileup with -q 40. We then plotted the allele frequency by subspecies (for the relevant two subspecies) for the alleles found in that genome assembly across each chromosome (using Pman2.1.3 coordinates), using 1 Mb windows. Through manual inspection of local ancestry plots, we identified four phasing errors, which aligned with the boundaries of four different contigs (Supplemental Figure S1). We fixed these phasing errors using bedtools getfasta to extract and exchange sequences. We then assigned chromosomes to subspecies based on the local ancestry analyses, and sorted chromosomes into subspecies genomes (since the draft assemblies were phased by chromosome only). Finally, we re-oriented chromosomes to match the direction of Pman2.1.3 chromosomes (e.g., Supplemental Figure S3), so that the four genome assemblies had consistent orientations across chromosomes; we took the reverse complement of reverse-oriented chromosomes using samtools fadix -i. We then recorded summary statistics for the final genome assemblies (Supplemental Table S1).

To visualize ancestry across the final genome assemblies (Figure 1B), we mapped whole-genome resequencing data (from NCBI PRJNA688305, PRJNA838595, PRJNA862503) from 15 individuals of each subspecies to the genome assembly for that subspecies and its pair (e.g., *rubidus* WGS data mapped to both *rubidus* and *gambelii* genomes), using 20M reads per individual and bwa -mem. We counted the number of mismatches per read using custom R scripts, averaging over 2 Mb windows by subspecies, and normalized the data by overall mean number of mismatches for a given chromosome.

### Repeat mining and annotation

We used RepeatModeler (version 2.0.2)^102^ to generate *de novo* transposable element libraries for each subspecies. We combined *de novo* repeat libraries with curated *P. maniculatus* TE models from^47^ and removed redundant models using CD-HIT-EST (version 4.8.1)^103^ with parameters -n 10 -c .8 -r 1 -i. Then, we combined these final *P. maniculatus* TE models with all rodent TE models available from DFAM^104^. We employed RepeatMasker (version 4.1.2)^105^ with parameters -pa 12 -excln -s -no_is -u -noisy -html -xm -a -xsmall to annotate interspersed and simple repeats in each genome using this combined set of interspersed repeat models as well as the previously identified *P. maniculatus* centromeric satellite sequence (NCBI accession: KX555281.1)^28^. Overlapping or redundant repeat annotations were resolved based on highest alignment scores through the RepeatMasker utility script RM2Bed.py^105^. RepeatMasker also masked repetitive sequences in each genome for downstream analysis. To identify segmental duplications for each subspecies, we used the masked version of each genome as input for BISER (version 1.4)^106^ with default parameters. We defined segmental duplications as duplications >1 kb in length with ≥90% identity when bases attributed to known high-copy-number repeats are ignored.

### Calling centromere positions and CpG methylation analysis

To annotate candidate centromere locations, we first extracted all centromeric satellite hits from our RepeatMasker output and merged all hits within 1 Mb of each other using bedtools merge (version 2.29.1). We considered all centromeric satellite arrays >100 kb as possible active centromeres. Some chromosomes displayed one distinct centromeric satellite array. However, others harbored multiple, distinct centromeric satellite arrays. While most centromeric sequences are highly methylated, active centromeres show signatures of DNA hypomethylation at the site of kinetochore assembly^29,30,35^. Thus, to resolve ambiguities and discern active centromeres, we predicted CpG methylation landscapes using PacBio HiFi kinetics data^107^. We did this using the following pipeline. First, we recalled PacBio HiFi consensus reads using pbccs (version 6.4.0) (https://ccs.how/) with the parameter --hifi-kinetics to include kinetics data for each read. Next, we employed jasmine (version 2.0.0) (https://github.com/pacificbiosciences/jasmine/) to predict per-site CpG methylation probabilities for all reads. Then, we aligned all reads annotated with CpG methylation probabilities to each respective genome assembly using pbmm2 (version 1.13.1) (https://github.com/PacificBiosciences/pbmm2) with the parameter --min-concordance-perc 99 to ensure only alignments for reads specific to the relevant haplotype were retained.

Pbmm2 is essentially a wrapper for minimap2 with additional utilities specifically for Pacbio data, including the ability to retain CpG methylation data in mapped bam output files. Using these alignments, we employed aligned_bam_to_cpg_scores (version 2.3.0) (https://github.com/PacificBiosciences/pb-CpG-tools) with the default model to obtain final CpG methylation predictions in bed format for each species. For each centromeric satellite array, we generated heatmaps displaying average CpG methylation probabilities across 5 kb windows using python. We normalized and adjusted CpG methylation probabilities to range from -1 to 1 to visually distinguish patterns of methylation more easily. Then, we manually called active centromeres using patterns of hypomethylation. We took a conservative approach, and marked chromosomes as inconclusive if more than one centromeric satellite array displayed evidence of a possible hypomethylated region or if a centromeric satellite array displayed noisy CpG methylation results. To visualize centromere structure, we generated all-by-all pairwise sequence identity plots for each centromere using StainedGlass (version 0.5)^108^ with default parameters. We used HiCAT (version 1.1.0)^54^ to predict HOR structure in all assembled centromeric arrays and plot HOR distributions.

### Alignment and structural variant calling

We called structural variants for *P. m. gambelii, P. m. rubidus,* and *P. m. nubiterrae* subspecies with respect to the *P. m. bairdii* genome assembly. We aligned *de novo* genome assemblies for *gambelii, rubidus and nubiterrae* to *bairdii* using minimap2 (version 2.21)^101^ with the setting -ax asm20 to allow for alignment of more divergent genomic regions. To call structural variants for each subspecies relative to *bairdii*, we used two complementary tools: SVIM-asm (version 1.0.3)^36^, which is sensitive to smaller structural variants, and SyRI (version 1.6.3)^37^, which uses alignment of syntenic regions to accurately detect larger structural rearrangements. Additionally, SyRI identifies balanced structural variants such as inversions from whole genome alignments with higher accuracy than other tools^37^. Both of these tools produced VCFs for each subspecies, which were merged by subspecies, and then across subspecies, using SURVIVOR (version 1.0.7)^109^ with parameters 500 1 1 1 0 50. Due to the challenges with calling structural variants in highly repetitive regions, we also filtered out SVs primarily composed of simple repeats^110^. After filtering, we were left with a combined set of 581,552 SVs, including 182,271 SVs and 15,529 SVs uniquely identified by SVIM-asm and SyRI respectively (Supplemental Figure S13). Inversions >1 Mb were validated manually using dot plots generated from whole-genome alignments from minimap2 with -cx asm5. In total, we identified 572,400 indels, 8,038 inversions, and 1,114 duplications across these four subspecies.

### Polarizing inversions

To polarize the megabase-scale inversions, we compared our *bairdii* genome assembly to the *Peromyscus leucopus* and the *Onychomys torridus* genomes (RefSeq accessions: GCF_004664715.2 and GCF_903995425.1 respectively), two outgroup species. We selected *P. leucopus* and *O. torridus* as outgroups for several reasons. First, both species are relatively closely related to *P. maniculatus* (<15 million years diverged). Second, the two species exhibit the same number of chromosomes as *P. maniculatus*. Third, the genomes of both species were produced with long-read sequencing technologies and display chromosome-level contiguity. We aligned our *P. m. bairdii* genome assembly to the two outgroup genomes using minimap2 with the setting -ax asm20. We called each *bairdii* allele as ancestral or derived based on the orientation of its alignment to *P. leucopus* and *O. torridus*. If the majority of the region corresponding to an inversion was collinear between *bairdii* and *P. leucopus* and *O. torridus*, we called the *bairdii* allele as ancestral, whereas if the majority of the region corresponding to an inversion was inverted in *P. leucopus* and *O. torridus* relative to *bairdii*, we called the *bairdii* allele as derived. In cases where alignments to *O. torridus* showed ambiguity, we called the ancestral allele using alignments to *P. leucopus*. We manually inspected alignments using dotplots to validate calls for all megabase-scale inversions as well as to resolve possible recurrent inversions.

### Inversion breakpoint analyses

We first tested for an enrichment of repeats at inversion breakpoints. To do so, we performed a permutation test for enrichment of repeats in breakpoint regions for five types of repeats: LINE retrotransposons, SINE retrotransposons, LTR retrotransposons, centromeric satellite repeats and segmental duplications. Specifically, we extracted the flanking regions of each inversion breakpoint using bedtools flank (version 2.29.1) and compared multiple metrics of repeat composition for these regions relative to expectations from randomization across 1,000 permutations using GAT (version 1.3.5)^111^. These metrics included the number of repeats overlapping inversion breakpoint regions, as well as the proportion of base pairs attributed to repeats in those regions. We performed these tests on our entire set of inversions for several flanking region sizes, including 100 bp, 200 bp, 1 kb, 10 kb and 500 kb. Since we hypothesized that smaller inversions (<1 Mb) and large inversions (>1 Mb) might show divergent origins, we also performed separate permutation tests for small and large inversions. All inversions and those those <1 Mb displayed enrichment (P<0.05) for SINEs, LINEs, LTRs and SDs in their breakpoint regions, whereas >1 Mb inversions displayed enrichment for SDs, centromeric repeats and LTRs.

Inversions commonly arise as a result of ectopic recombination between repetitive elements^16^. To search for evidence of repeat-mediated inversions, we intersected inversion breakpoint regions with our TE and SD annotations, and performed a similar procedure as described in^19^. Briefly, we searched for homozygous TEs in the 200 bp regions flanking each inversion. Due to the size of SDs and the challenges associated with localizing inversion breakpoints in SD-enriched regions, we extended our search for SDs at inversion breakpoints by 50 kb in each direction for inversions >100 kb and <1 Mb, and by 500 kb for inversions >1 Mb. We called repeat-mediated inversions based on the presence of TEs from the same family at both breakpoints, or flanking SDs at both breakpoints (noting that we did not require SD homology). To investigate the relationship between inversion length and flanking SD length, we performed linear regressions comparing inversion length to average flanking SD length (Figure 4J) as well as maximum flanking SD length (Supplemental Figure S11). Both correlations were highly significant (Kendall’s Tau = 0.216 and 0.231, P<0.00001 and P<0.00001 respectively). We considered all inversions flanked by SDs at both breakpoints in these analyses. To visualize SD-enriched inversion breakpoints, we created dotplots of the breakpoint regions. Self-v-self alignments by chromosome were performed for the *P. m. bairdii* genome using nucmer (mummer version 4.0.0) with –maxmatch –nosimplify -l 50 -c 100. Alignments >1 kb were plotted using R.

### Analyzing SD and TE distributions and the origins of SDs

To investigate whether TEs are associated with SD formation, we compared Kernel Density Estimates (derived using python) for SD divergences and TE divergences from consensus sequences^105,106^. To search for patterns consistent with SD origins from TE-mediated ectopic recombination, we identified SDs flanked by older, related TEs. Then, we compared the number of SDs exhibiting this pattern to expectations from random resampling across 1,000 permutations. We performed a linear regression for SD divergence and average associated TE divergence using python. Tests for enrichment of different TE classes close to centromeres were performed using the same method as for SDs. “Young TEs” were called using a maximum Kimura Divergence from the consensus of 10%. Analyses performed with other cutoffs produced the same results, concordant with strong correlations between TE Kimura Divergence and distance from the nearest centromeric satellite array on continuous scales.

### Centromeric satellite-mediated inversions

To understand the role of centromeric satellite arrays in the evolution of inversions, we analyzed the nine mega-base scale inversions with centromeric satellites within 1 Mb of both breakpoints (recurrent inversions and nested inversion regions were excluded). We identified the ancestral haplotype for each of these inversions (see “Polarizing inversions” section) and evaluated the orientation of the centromeric satellite repeats at these ancestral inversion breakpoints using annotations from repeatmasker, and plotted satellite locations and orientations using python.

## Acknowledgements

The authors thank Russell Corbett-Detig, Bohao Fang, Andreas Kautt, Amanda Larracuente, Peter Sudmant, and Tianzhu Xiong for helpful discussions and feedback on this manuscript. The genomes were assembled and sequenced by Cantata Bio (Dovetail Genomics), Scotts Valley, CA. The computations for this work were run on the FASRC Cannon cluster supported by the FAS Division of Science Research Computing Group at Harvard University. O.S.H. was supported by a National Science Foundation Graduate Research Fellowship. This work was supported by the Howard Hughes Medical Institute.

## Author contributions

O.S.H., L.G. and H.E.H. conceived of and designed the research. L.G. and O.S.H. performed the research and analyses. L.G., O.S.H. and H.E.H. wrote the manuscript.

---

**Supplemental Figure S1:**
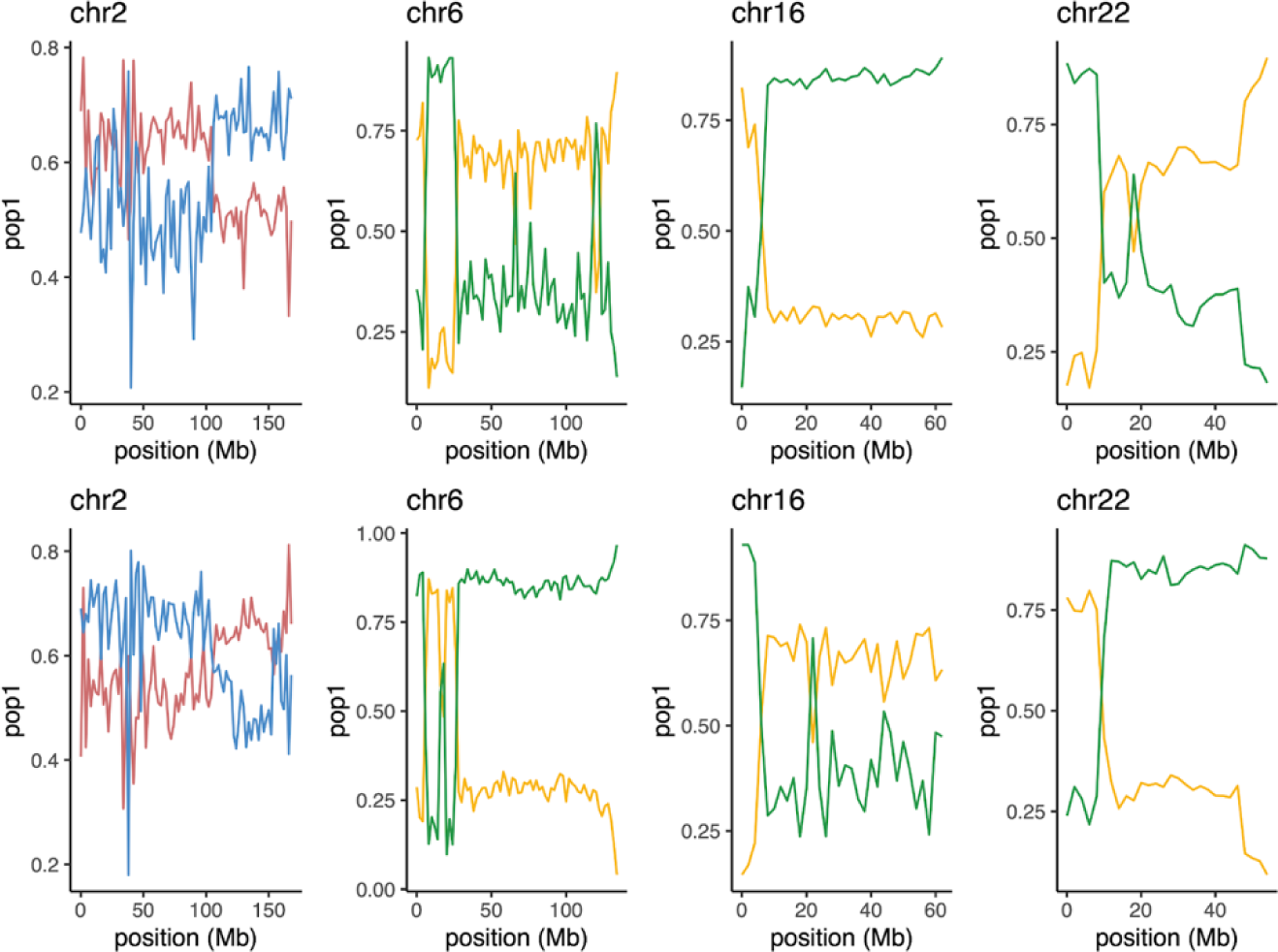
Phasing errors identified during quality control of *de novo* genome assemblies. A total of four phasing errors were identified by analyzing ancestry of the phased contigs. Contigs from each genome were mapped to the Pman2.1.3 reference genome chromosomes (x-axis), and ancestry was determined using short-read whole-genome sequencing data from each subspecies (see Methods). Average allele frequencies for ancestry-specific SNPs are shown in 2 Mb windows. Colors by subspecies: red=*rubidus*; blue=*gambelii*; yellow=*bairdii*; green=*nubiterrae*.

**Supplemental Figure S2:**
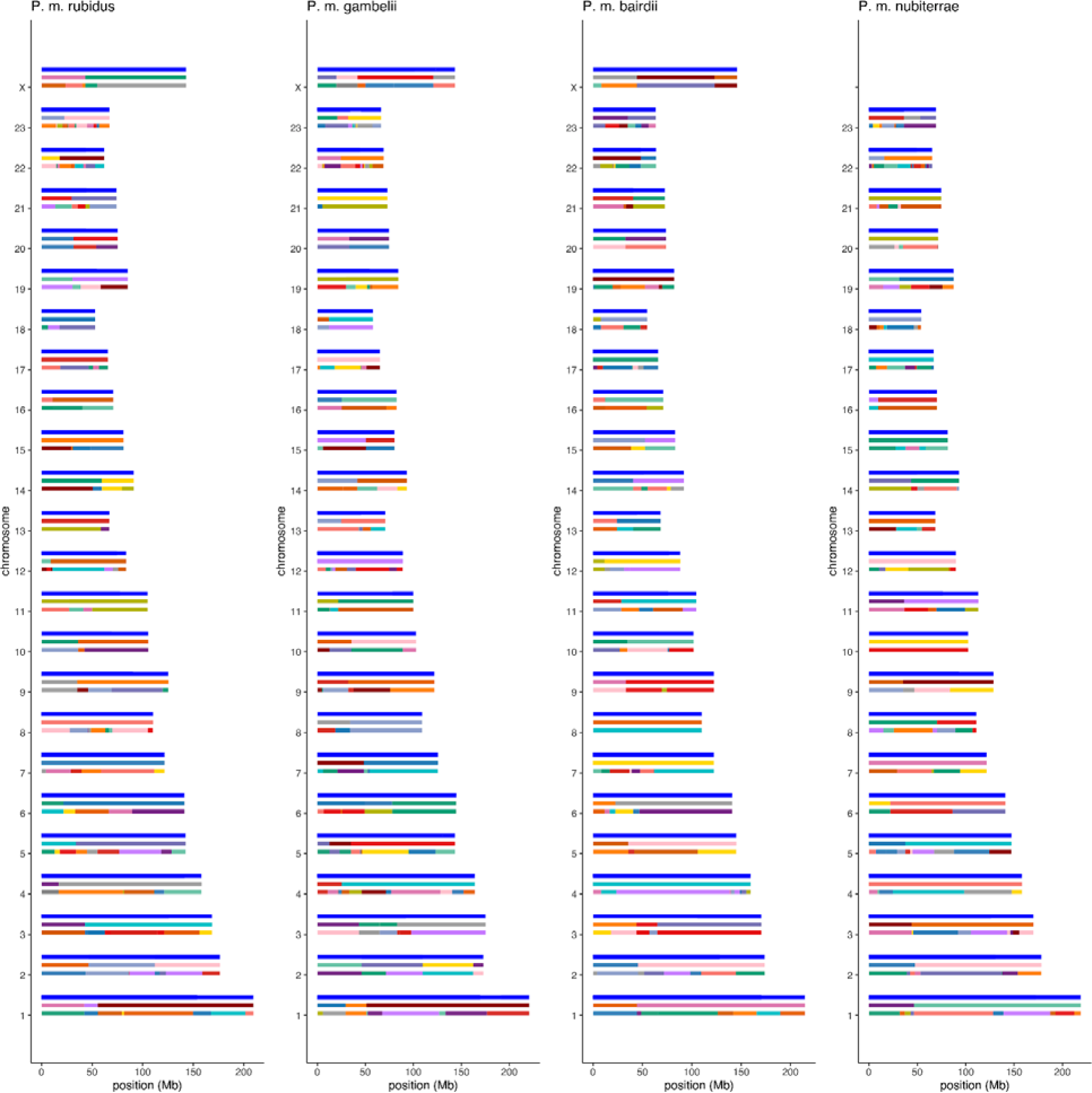
Assembly of four *de novo* genomes. Contig-level genome assemblies were created with *hifi-asm*, with positions and number of contigs shown for each chromosome (bottom row). Contigs were scaffolded using *HiRise*, with positions and number of *HiRise* scaffolds shown for each chromosome (middle row). Full chromosomes (top row) were assembled through manually joining *HiRise* scaffolds using juicer. Subspecies labels are provided for each genome. Note that a male haplotype for *P. m. nubiterrae* was sequenced, explaining the missing X-chromosome for *P. m. nubiterrae*.

**Supplemental Figure S3:**
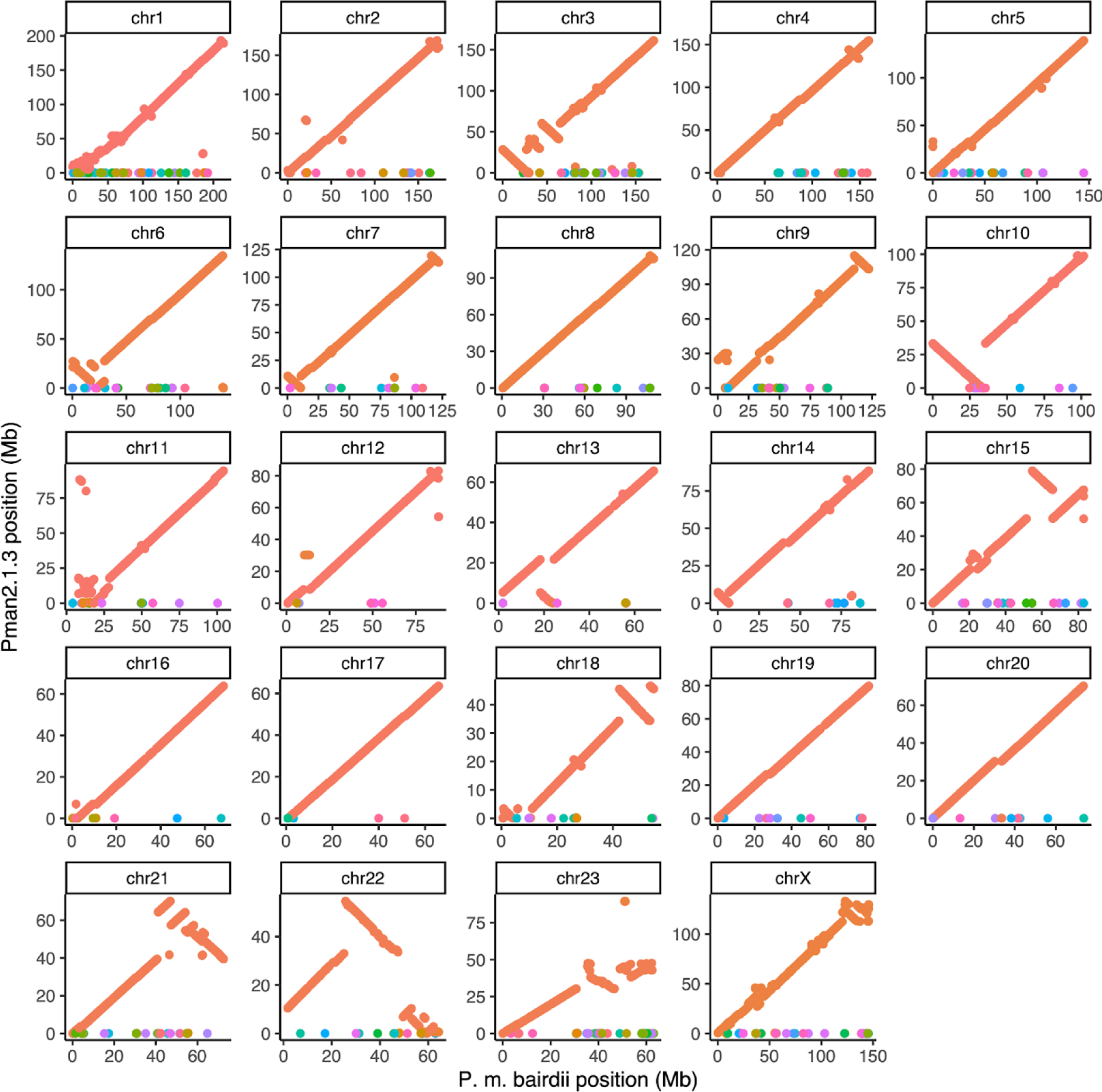
Alignments of *de novo* genome assembly for *P. m. bairdii* with Pman2.1.3 reference genome, shown for the 24 major scaffolds in the *de novo* assembly. All alignments >1 kb from minimap2 alignments are plotted as dots and colored by scaffold in the Pman2.1.3 assembly, with Pman2.1.3 chromosomes colored in orange.

**Supplemental Figure S4:**
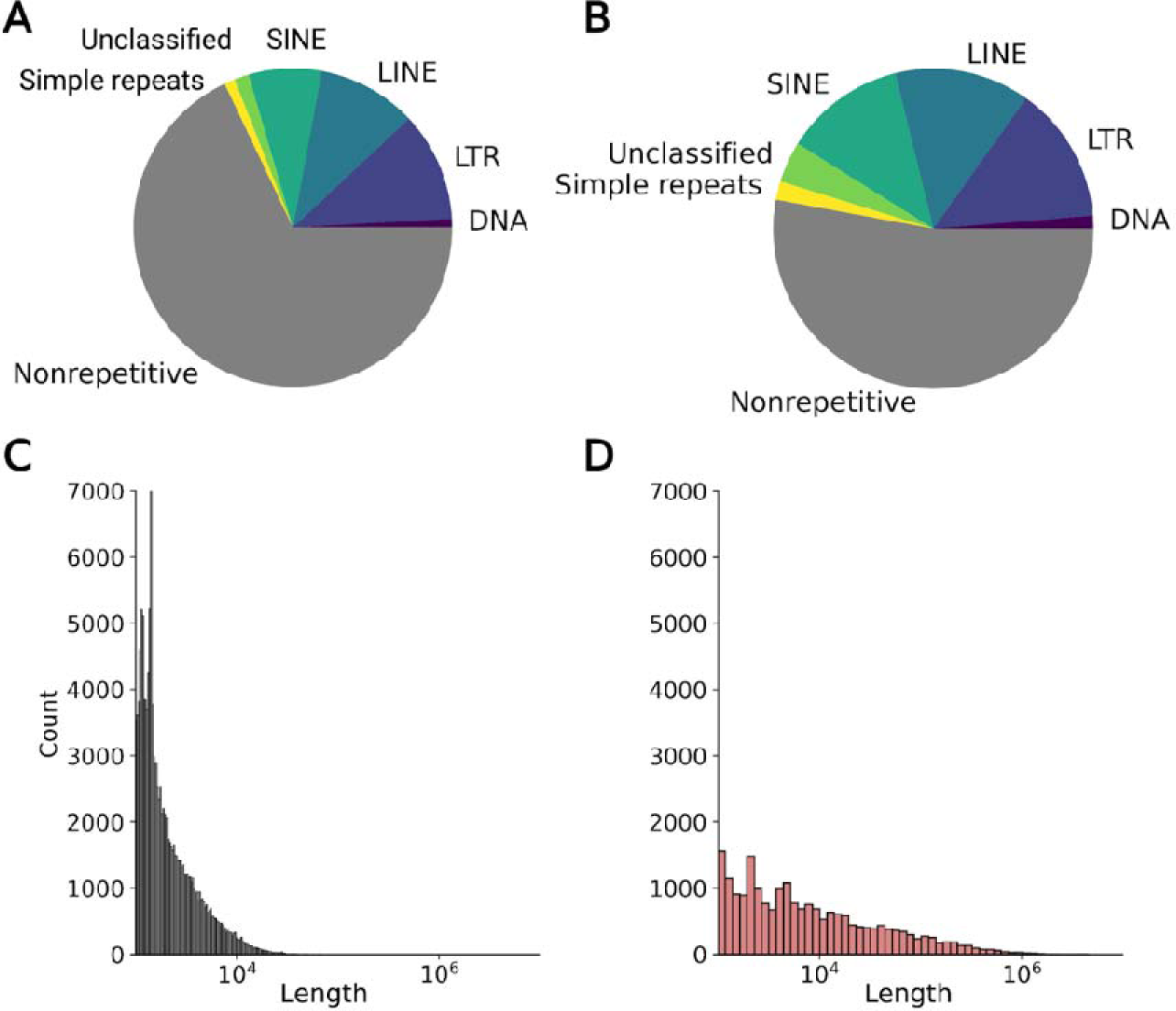
(**A**-**B**) Repeat landscapes for (**A**) the previous short-read-based *P. maniculatus bairdii* (RefSeq: GCF_003704035.1) reference assembly and (**B**) the new *P. maniculatus bairdii* assembly presented in this work. The new assembly shows a 13% increase in interspersed and simple repeat occupancy relative to the old assembly, demonstrating a significant increase in repeat representation and assembly quality. (**C**-**D**) Segmental duplication length distributions for (C) the previous *P. m. bairdii* assembly and (D) the new *P. m. bairdii* assembly. The SD length distribution is shifted to the right in the new assembly, indicating higher contiguity. X-axes are logscaled.

**Supplemental Figure S5:**
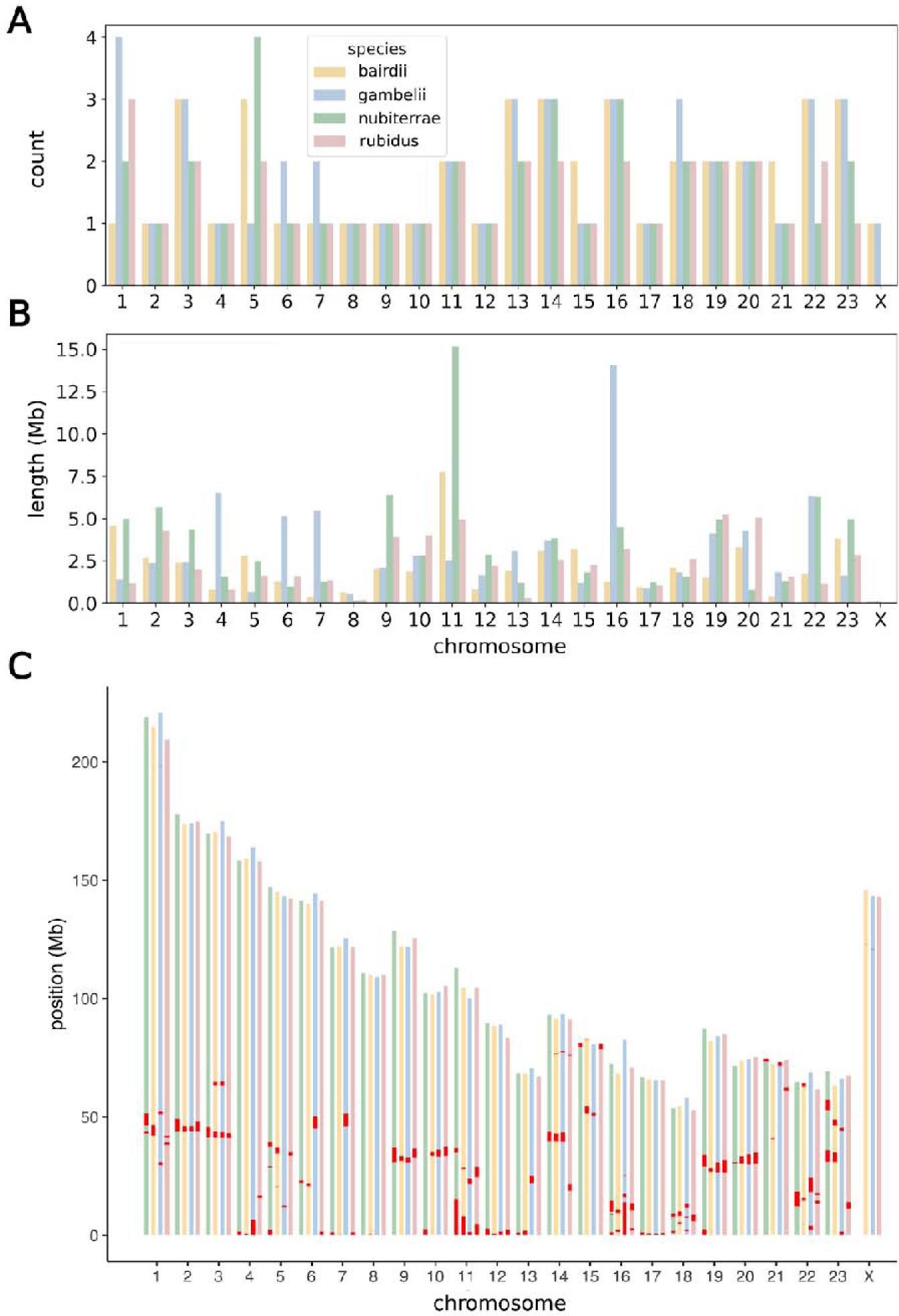
(**A**) Number of interspersed centromeric satellite arrays (>1 kb) for each chromosome across subspecies. (**B**) Total aggregate length of sequence occupied by centromeric satellite arrays for each chromosome in each subspecies. (**C**) Genomic position for each centromeric satellite array (red) on each chromosome across subspecies. Subspecies colors correspond to legend in (A).

**Supplemental Figure S6:**
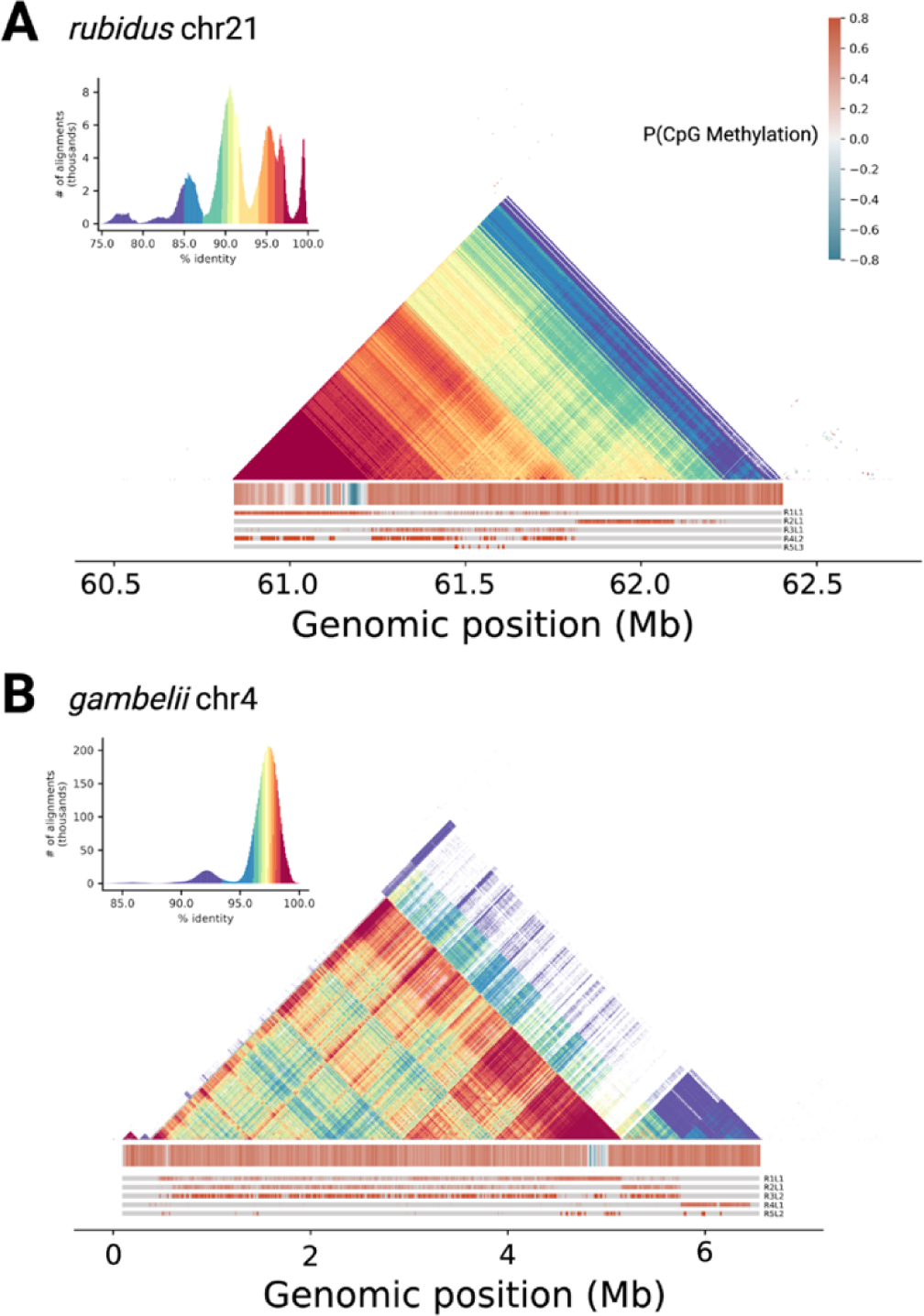
Additional examples of centromere structure for (**A**) *rubidus* chr21 and (**B**) *gambelii* chr4. From top to bottom in each panel: stained glass plots showing self-vs-self percent identity calculated across 5 kb windows, heatmaps displaying normalized probability of CpG methylation calculated across 5 kb windows, and heatmaps showing positions of the top 5 most common centromeric repeats for each centromere.

**Supplemental Figure S7:**
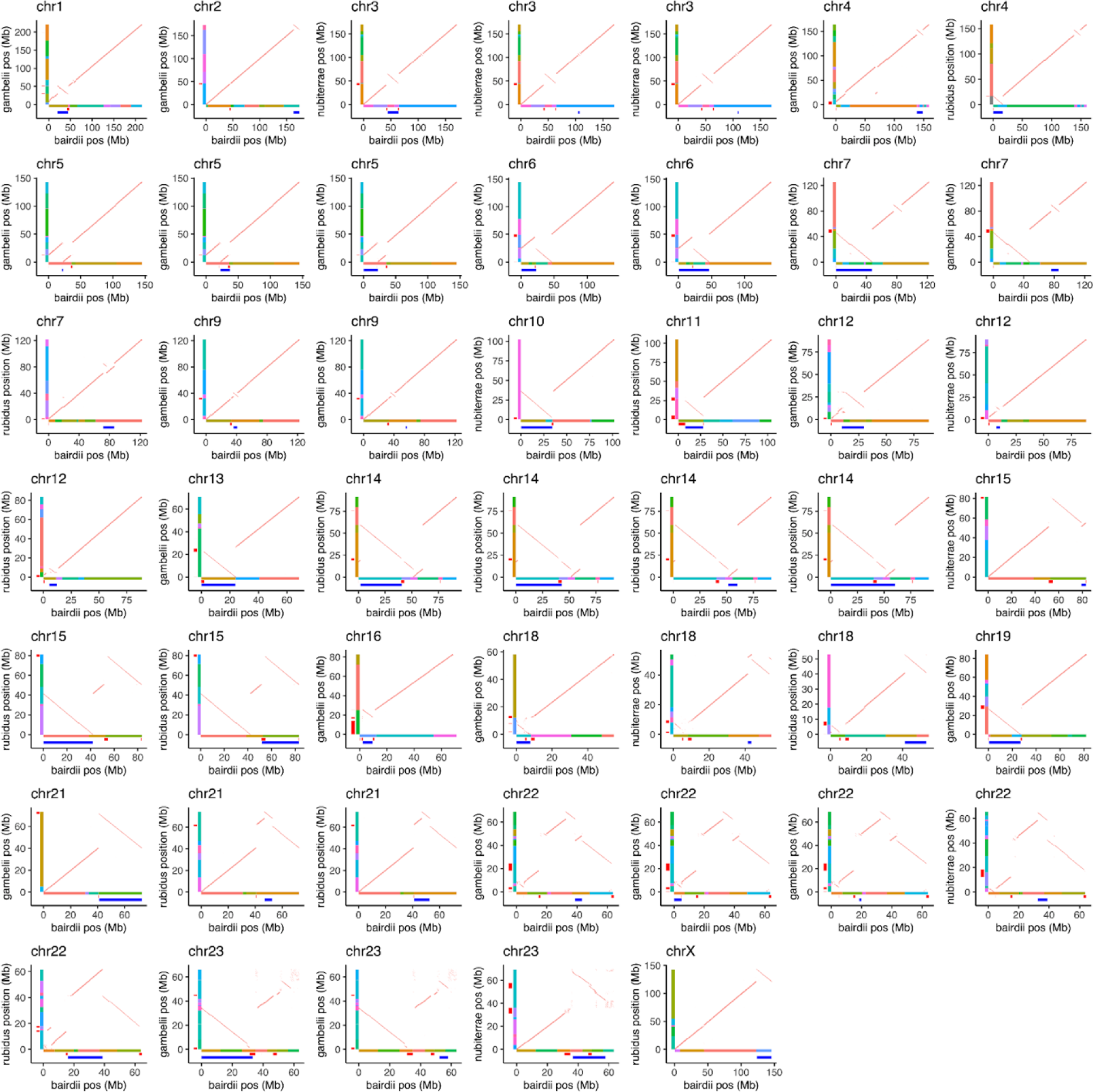
Alignments of *de novo* genome assemblies showing inversions. For each identified inversion, alignments between *P. m. bairdii* genome (x-axis) and relevant subspecies (y-axis) genome is shown in orange for the entire chromosome. Contigs from the initial *hifi-asm* assemblies are shown as colored rectangles, with locations of centromere satellite arrays shown as red rectangles, and identified inversions shown as blue rectangles.

**Supplemental Figure S8:**
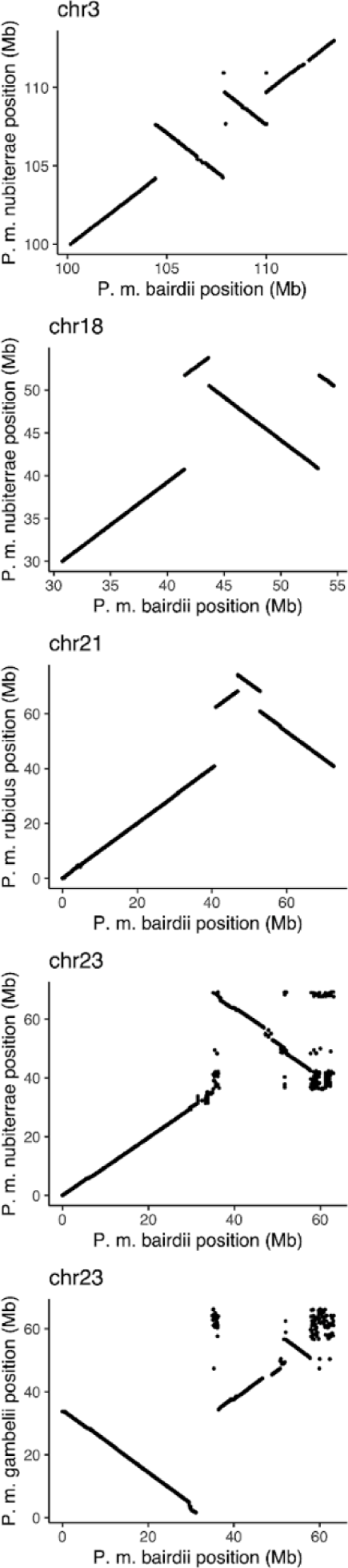
Example of recurrent inversion breakpoints. Alignments (>10 kb) between *P. m. bairdii* and relevant subspecies are shown for chromosomes harboring recurrent inversion breakpoints.

**Supplemental Figure S9:**
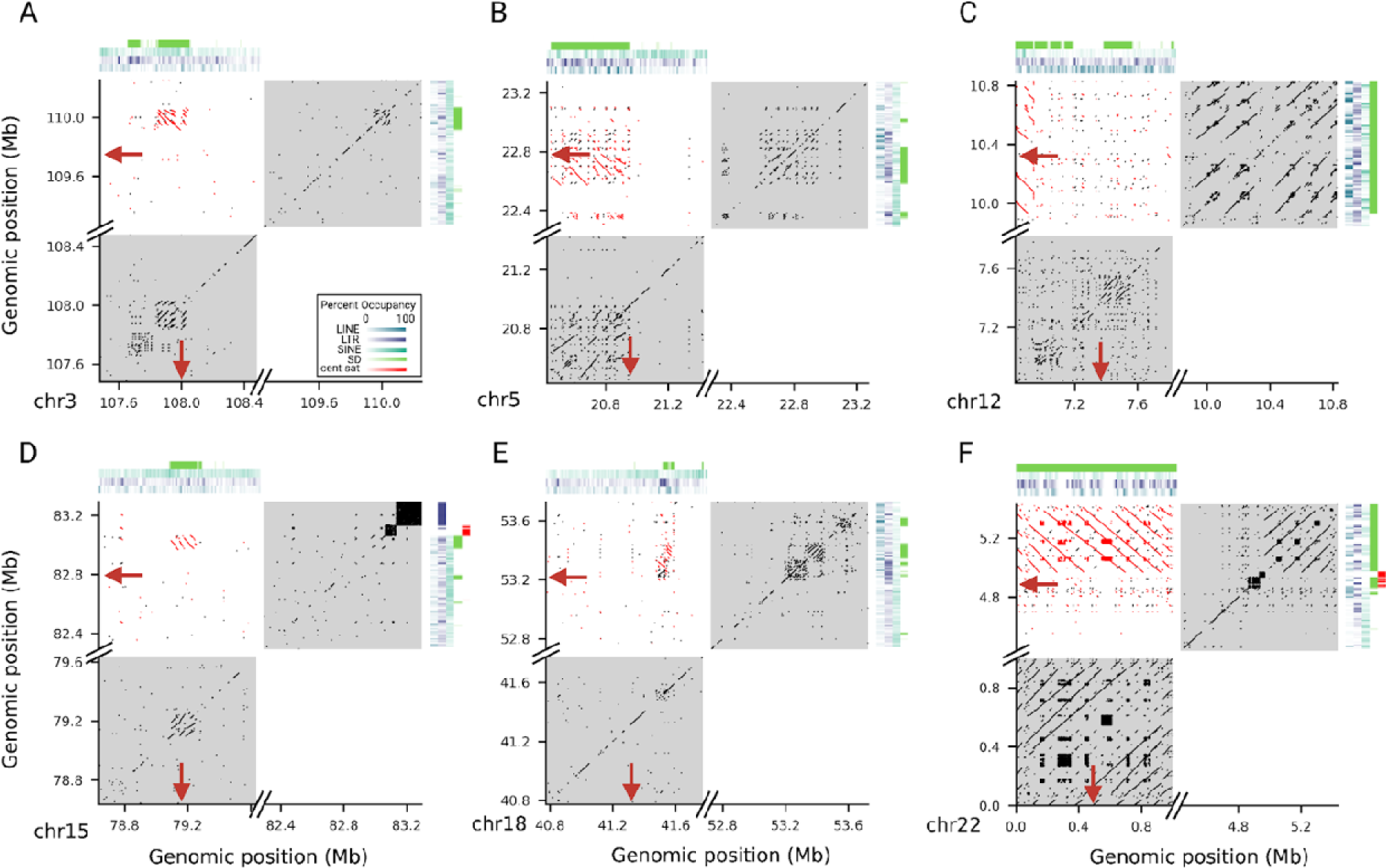
(**A**-**F**) Examples of inversion breakpoints near large inverted segmental duplications. Dotplots show self-v-self as well as breakpoint-v-breakpoint alignments for inversion breakpoints in *P. maniculatus bairdii.* Inverted alignments are plotted in red and collinear alignments are plotted in black. Self-v-self alignments are highlighted with gray boxes. Inversion breakpoints are annotated with red arrows. Alignments within 500 kb of breakpoints are shown. Only alignments >100 bp were included. Heatmaps show repeat occupancy for various repeat types calculated across 10 kb windows. The key in (A) corresponds to heatmap colors.

**Supplemental Figure S10:**
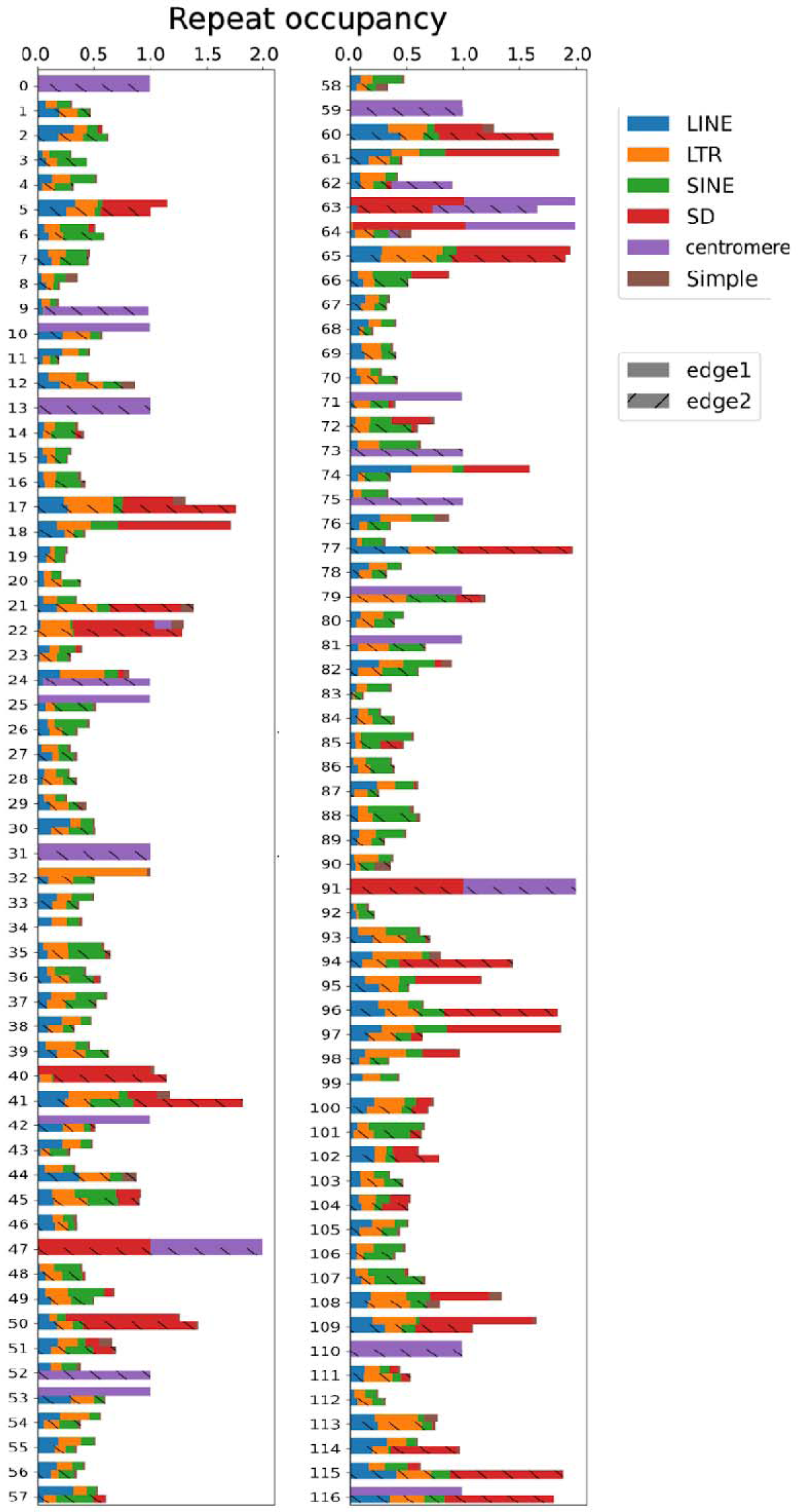
Repeat landscapes for 100 kb edges of neighboring contigs for all 117 contig scaffolding events in the *P. maniculatus bairdii* genome. Considered repeats include LINE, SINE, and LTR retrotransposons, centromeric satellite, simple repeats and SDs. Repeat occupancy is calculated by dividing the number of bases occupied by a given repeat by 100 kb. Usually, occupancy should add to 1. However, SDs often contain other interspersed repeats and thus introduce redundancy to occupancy calls. As a result, the maximum occupancy in this case is 2.

**Supplemental Figure S11:**
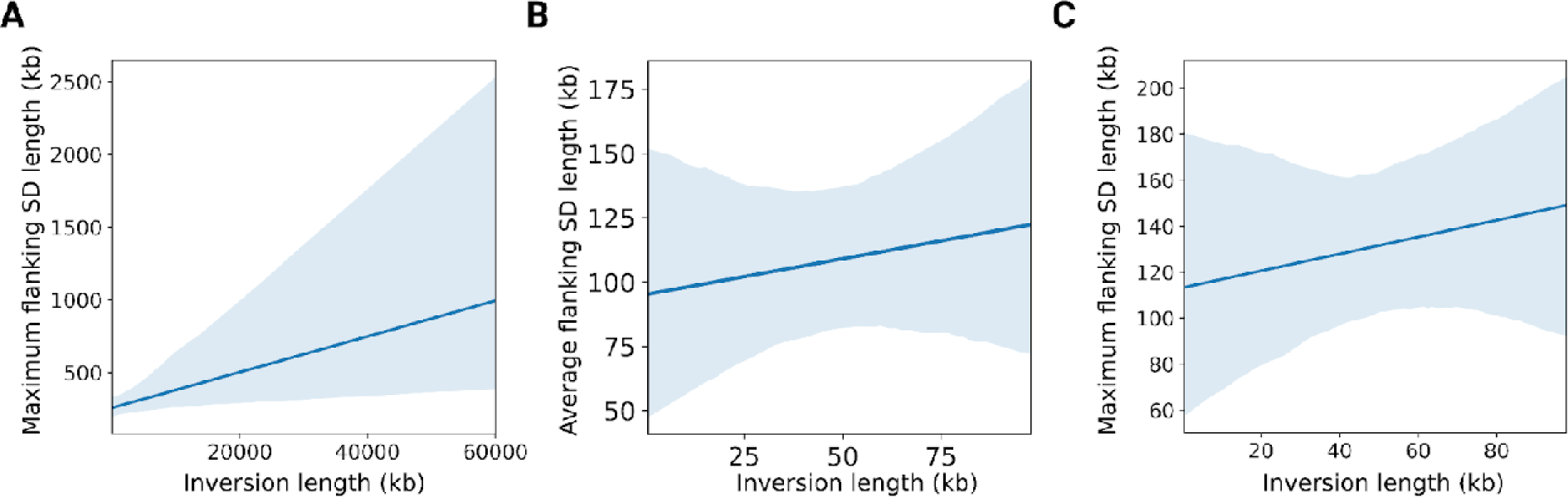
(**A**) Linear regression with a “robust” model accounting for outlier effects^112^ for maximum flanking SD length as a function of associated inversion length (Kendall’s tau = 0.231, P<0.0001). Although both the regression model used here and Kendall’s nonparametric correlation metrics should be robust to outlier effects, we also performed linear regressions comparing (**B**) average and (**C**) maximum flanking SD length to inversion length for inversions <100 kb, to confirm that correlations are not merely driven by larger inversions (Kendall’s tau = 0.172, 0.185 and P= 0.0088,0.0051 respectively).

**Supplemental Figure S12:**
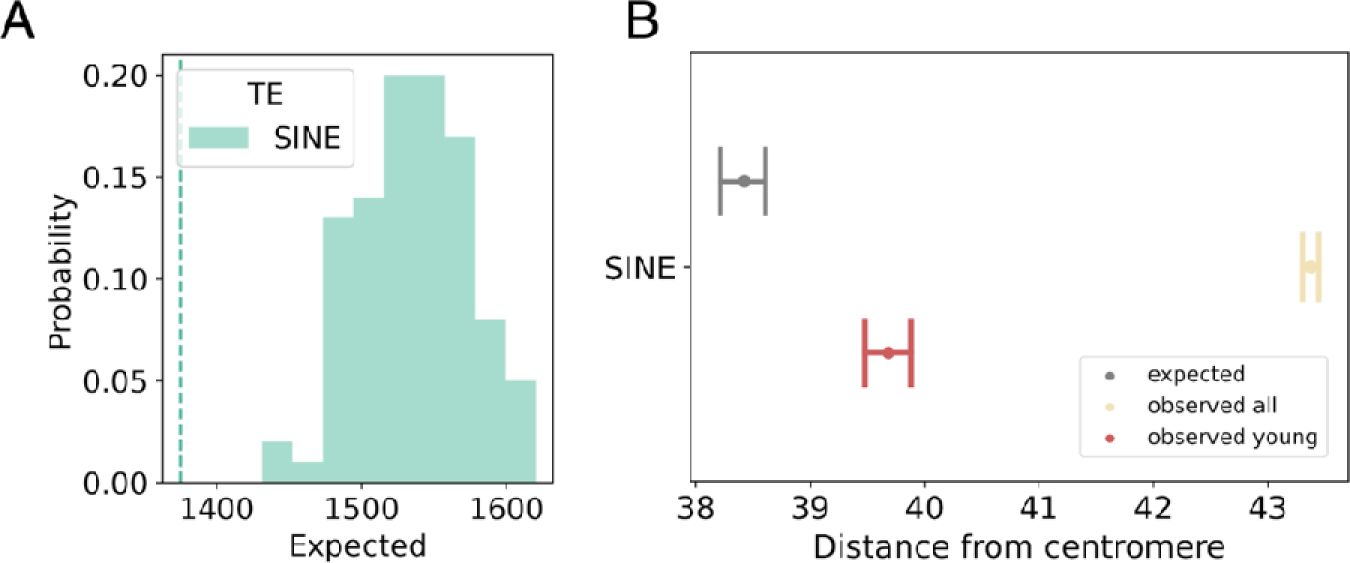
(**A**) Number of SDs showing patterns consistent with origins from SINE-mediated ectopic recombination (see Figure 5C) compared to expectations from random resampling. Histograms show expected distributions and dotted lines denote observed values (P>0.99). (**B**) Observed distributions of distances from the nearest centromeric repeat array compared to expectations from random resampling for SINEs.

**Supplemental Figure S13:**
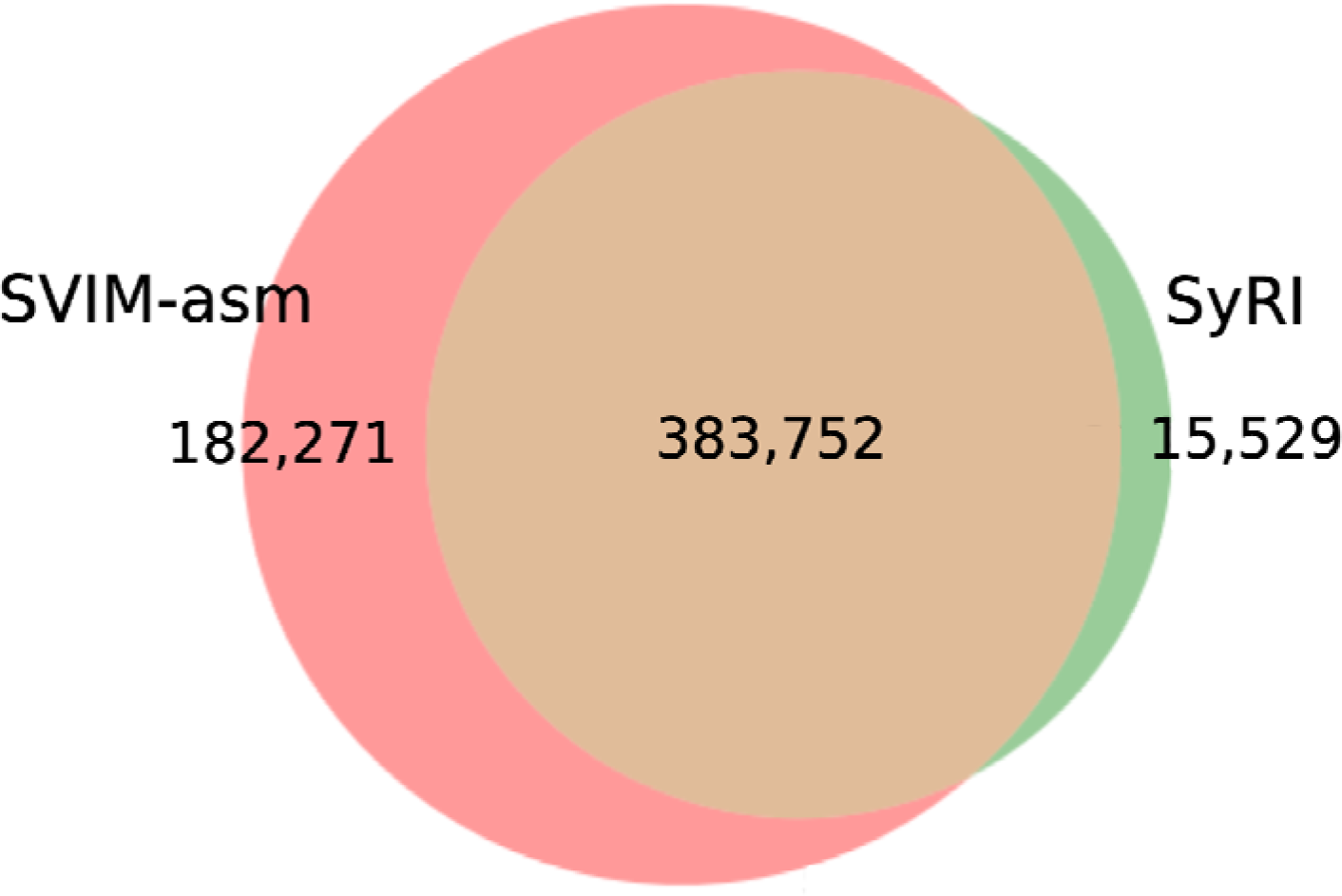
Venn diagram showing the number of SVs unique to each structural variant caller as well as the number of SVs supported by both callers.

## Notes

### Competing Interest Statement

The authors have declared no competing interest.

## References

1. Hecker, N. & Hiller, M. A genome alignment of 120 mammals highlights ultraconserved element variability and placenta-associated enhancers. Gigascience 9, (2020).

2. Lindblad-Toh, K. et al. A high-resolution map of human evolutionary constraint using 29 mammals. Nature 478, 476–482 (2011).

3. Sturtevant, A. H. A Case of rearrangement of genes in *Drosophila*. Proceedings of the National Academy of Sciences 7, 235–237 (1921).

4. Cross, J. C. A comparative study of the chromosomes of rodents. J. Morphol. 52, 373–401 (1931).

5. White, M. J. D. Chromosomal rearrangements and speciation in animals. Annu. Rev. Genet. 3, 75–98 (1969).

6. Stebbins, G. L. Chromosomal variation and evolution. Science 152, 1463–1469 (1966).

7. Chu, E. H. & Bender, M. A. Chromosome cytology and evolution in primates. Science 133, 1399– 1405 (1961).

8. Fan, Y., Linardopoulou, E., Friedman, C., Williams, E. & Trask, B. J. Genomic structure and evolution of the ancestral chromosome fusion site in 2q13-2q14.1 and paralogous regions on other human chromosomes. Genome Res. 12, 1651–1662 (2002).

9. J L Patton, A. & Sherwood, S. W. Chromosome evolution and speciation in rodents. Annual Review of Ecology and Systematics 14, 139–158 (1983).

10. White, M. J. D. Animal Cytology and Evolution. (Cambridge University Press, 1955).

11. Coyne, J. A., Meyers, W., Crittenden, A. P. & Sniegowski, P. The fertility effects of pericentric inversions in *Drosophila melanogaster*. Genetics 134, 487–496 (1993).

12. Mérot, C., Oomen, R. A., Tigano, A. & Wellenreuther, M. A roadmap for understanding the evolutionary significance of structural genomic variation. Trends Ecol. Evol. 35, 561–572 (2020).

13. Wellenreuther, M. & Bernatchez, L. Eco-evolutionary genomics of chromosomal inversions. Trends Ecol. Evol. 33, 427–440 (2018).

14. Corbett-Detig, R. B. et al. Fine-mapping complex inversion breakpoints and investigating somatic pairing in the *Anopheles gambiae* species complex using proximity-ligation sequencing. Genetics 213, 1495–1511 (2019).

15. Villoutreix, R. et al. Inversion breakpoints and the evolution of supergenes. Mol. Ecol. 30, 2738– 2755 (2021).

16. Porubsky, D. et al. Recurrent inversion toggling and great ape genome evolution. Nat. Genet. 52, 849–858 (2020).

17. Ranz, J. M. et al. Principles of genome evolution in the *Drosophila melanogaster* species group. PLoS Biol. 5, e152 (2007).

18. Corbett-Detig, R. B. & Hartl, D. L. Population genomics of inversion polymorphisms in *Drosophila melanogaster*. PLoS Genet. 8, e1003056 (2012).

19. Porubsky, D. et al. Recurrent inversion polymorphisms in humans associate with genetic instability and genomic disorders. Cell 185, 1986–2005.e26 (2022).

20. Porubsky, D. et al. Inversion polymorphism in a complete human genome assembly. Genome Biol. 24, 100 (2023).

21. da Silva, V. H. et al. The genomic complexity of a large inversion in great tits. Genome Biol. Evol. 11, 1870–1881 (2019).

22. Fang, Z. et al. Megabase-scale inversion polymorphism in the wild ancestor of maize. Genetics 191, 883–894 (2012).

23. Sanchez-Donoso, I. et al. Massive genome inversion drives coexistence of divergent morphs in common quails. Curr. Biol. 32, 462–469.e6 (2022).

24. Bradshaw, W. N. & Hsu, T. C. Chromosomes of *Peromyscus* (Rodentia, cricetidae). Cytogenet. Genome Res. 11, 436–451 (1972).

25. Sparkes, R. S. & Arakaki, D. T. Intrasubspecific and intersubspecific chromosomal polymorphism in *Peromyscus maniculatus* (deer mouse). Cytogenet. Genome Res. 5, 411–418 (1966).

26. Harringmeyer, O. S. & Hoekstra, H. E. Chromosomal inversion polymorphisms shape the genomic landscape of deer mice. Nat Ecol Evol 6, 1965–1979 (2022).

27. Hager, E. R. et al. A chromosomal inversion contributes to divergence in multiple traits between deer mouse ecotypes. Science 377, 399–405 (2022).

28. Smalec, B. M., Heider, T. N., Flynn, B. L. & O’Neill, R. J. A centromere satellite concomitant with extensive karyotypic diversity across the *Peromyscus* genus defies predictions of molecular drive. Chromosome Res. 27, 237–252 (2019).

29. Logsdon, G. A. & Eichler, E. E. The dynamic structure and rapid evolution of human centromeric satellite dna. Genes 14, (2022).

30. Altemose, N. et al. Complete genomic and epigenetic maps of human centromeres. Science 376, eabl4178 (2022).

31. Talbert, P. B. & Henikoff, S. What makes a centromere? Exp. Cell Res. 389, 111895 (2020).

32. Barra, V. & Fachinetti, D. The dark side of centromeres: types, causes and consequences of structural abnormalities implicating centromeric DNA. Nat. Commun. 9, 4340 (2018).

33. Stimpson, K. M., Matheny, J. E. & Sullivan, B. A. Dicentric chromosomes: unique models to study centromere function and inactivation. Chromosome Res. 20, 595–605 (2012).

34. Gascoigne, K. E. & Cheeseman, I. M. Induced dicentric chromosome formation promotes genomic rearrangements and tumorigenesis. Chromosome Res. 21, 407–418 (2013).

35. Ichikawa, K. et al. Centromere evolution and CpG methylation during vertebrate speciation. Nat. Commun. 8, 1833 (2017).

36. Heller, D. & Vingron, M. SVIM-asm: structural variant detection from haploid and diploid genome assemblies. Bioinformatics 36, 5519–5521 (2021).

37. Goel, M., Sun, H., Jiao, W.-B. & Schneeberger, K. SyRI: finding genomic rearrangements and local sequence differences from whole-genome assemblies. Genome Biol. 20, 277 (2019).

38. Cáceres, M., Sullivan, R. T. & Thomas, J. W. A recurrent inversion on the eutherian X chromosome. Proc. Natl. Acad. Sci. U. S. A. 104, 18571–18576 (2007).

39. Krimbas, C. B. & Powell, J. R. Drosophila Inversion Polymorphism. (CRC Press, 1992).

40. Cáceres, M., Ranz, J. M., Barbadilla, A., Long, M. & Ruiz, A. Generation of a widespread *Drosophila* inversion by a transposable element. Science 285, 415–418 (1999).

41. Lim, J. K. & Simmons, M. J. Gross chromosome rearrangements mediated by transposable elements in *Drosophila melanogaster*. Bioessays 16, 269–275 (1994).

42. Lonnig, W.-E. & Saedler, H. Chromosome rearrangements and transposable elements. Annu. Rev. Genet. 36, 389–410 (2002).

43. Delprat, A., Negre, B., Puig, M. & Ruiz, A. The transposon Galileo generates natural chromosomal inversions in *Drosophila* by ectopic recombination. PLoS One 4, e7883 (2009).

44. Bailey, J. A., Liu, G. & Eichler, E. E. An Alu transposition model for the origin and expansion of human segmental duplications. Am. J. Hum. Genet. 73, 823–834 (2003).

45. Kazazian, H. H., Jr. Mobile elements: drivers of genome evolution. Science 303, 1626–1632 (2004).

46. Nei, M. & Kumar, S. Molecular Evolution and Phylogenetics. (Oxford University Press, 2000).

47. Gozashti, L., Feschotte, C. & Hoekstra, H. E. Transposable element interactions shape the ecology of the deer mouse genome. Mol. Biol. Evol. 40, (2023).

48. Dittwald, P. et al. NAHR-mediated copy-number variants in a clinical population: mechanistic insights into both genomic disorders and Mendelizing traits. Genome Res. 23, 1395–1409 (2013).

49. Campos-Sánchez, R., Cremona, M. A., Pini, A., Chiaromonte, F. & Makova, K. D. Integration and fixation preferences of human and mouse endogenous retroviruses uncovered with functional data analysis. PLoS Comput. Biol. 12, e1004956 (2016).

50. Graham, T. & Boissinot, S. The genomic distribution of L1 elements: the role of insertion bias and natural selection. J. Biomed. Biotechnol. 2006, 75327 (2006).

51. Medstrand, P., van de Lagemaat, L. N. & Mager, D. L. Retroelement distributions in the human genome: variations associated with age and proximity to genes. Genome Res. 12, 1483–1495 (2002).

52. Schubert, I. What is behind ‘centromere repositioning’? Chromosoma 127, 229–234 (2018).

53. Anlaş, Ö. et al. Dicentric recombinant chromosome 18 due to maternal paracentric inversion analyzed by array CGH. Mol. Syndromol. 14, 246–253 (2023).

54. Gao, S. et al. HiCAT: a tool for automatic annotation of centromere structure. Genome Biol. 24, 58 (2023).

55. Melters, D. P. et al. Comparative analysis of tandem repeats from hundreds of species reveals unique insights into centromere evolution. Genome Biol. 14, R10 (2013).

56. Kirkpatrick, M. How and why chromosome inversions evolve. PLoS Biol. 8, (2010).

57. Faria, R., Johannesson, K., Butlin, R. K. & Westram, A. M. Evolving inversions. Trends Ecol. Evol. 34, 239–248 (2019).

58. Hoffmann, A. A. & Rieseberg, L. H. Revisiting the impact of inversions in evolution: from population genetic markers to drivers of adaptive shifts and speciation? Annu. Rev. Ecol. Evol. Syst. 39, 21–42 (2008).

59. Kent, T. V., Uzunović, J. & Wright, S. I. Coevolution between transposable elements and recombination. Philos. Trans. R. Soc. Lond. B Biol. Sci. 372, (2017).

60. Ling, A. & Cordaux, R. Insertion sequence inversions mediated by ectopic recombination between terminal inverted repeats. PLoS One 5, e15654 (2010).

61. Balachandran, P. et al. Transposable element-mediated rearrangements are prevalent in human genomes. Nat. Commun. 13, 7115 (2022).

62. Zhang, J. et al. Alternative Ac/Ds transposition induces major chromosomal rearrangements in maize. Genes Dev. 23, 755–765 (2009).

63. Klein, S. J. & O’Neill, R. J. Transposable elements: genome innovation, chromosome diversity, and centromere conflict. Chromosome Res. 26, 5–23 (2018).

64. Liu, P. et al. Frequency of nonallelic homologous recombination is correlated with length of homology: evidence that ectopic synapsis precedes ectopic crossing-over. Am. J. Hum. Genet. 89, 580–588 (2011).

65. Petrov, D. A., Aminetzach, Y. T., Davis, J. C., Bensasson, D. & Hirsh, A. E. Size matters: non-LTR retrotransposable elements and ectopic recombination in *Drosophila*. Mol. Biol. Evol. 20, 880–892 (2003).

66. De Coster, W., Weissensteiner, M. H. & Sedlazeck, F. J. Towards population-scale long-read sequencing. Nat. Rev. Genet. 22, 572–587 (2021).

67. Lupski, J. R. Genomic disorders: structural features of the genome can lead to DNA rearrangements and human disease traits. Trends Genet. 14, 417–422 (1998).

68. Stankiewicz, P. & Lupski, J. R. Genome architecture, rearrangements and genomic disorders. Trends Genet. 18, 74–82 (2002).

69. Bailey, J. A. & Eichler, E. E. Primate segmental duplications: crucibles of evolution, diversity and disease. Nat. Rev. Genet. 7, 552–564 (2006).

70. Rudd, M. K. & Willard, H. F. Analysis of the centromeric regions of the human genome assembly. Trends Genet. 20, 529–533 (2004).

71. Rocchi, M., Archidiacono, N., Schempp, W., Capozzi, O. & Stanyon, R. Centromere repositioning in mammals. Heredity 108, 59–67 (2012).

72. Šatović-Vukšić, E. & Plohl, M. Satellite DNAs-from localized to highly dispersed genome components. Genes 14, (2023).

73. Montefalcone, G., Tempesta, S., Rocchi, M. & Archidiacono, N. Centromere repositioning. Genome Res. 9, 1184–1188 (1999).

74. Emanuel, B. S. & Shaikh, T. H. Segmental duplications: an ‘expanding’ role in genomic instability and disease. Nat. Rev. Genet. 2, 791–800 (2001).

75. Charlesworth, B., Sniegowski, P. & Stephan, W. The evolutionary dynamics of repetitive DNA in eukaryotes. Nature 371, 215–220 (1994).

76. Walsh, J. B. Persistence of tandem arrays: implications for satellite and simple-sequence DNAs. Genetics 115, 553–567 (1987).

77. Smith, G. P. Evolution of repeated DNA sequences by unequal crossover. Science 191, 528–535 (1976).

78. Garrido-Ramos, M. A. Satellite DNA: an evolving topic. Genes 8, (2017).

79. Dover, G. Molecular drive: a cohesive mode of species evolution. Nature 299, 111–117 (1982).

80. Plohl, M., Meštrović, N. & Mravinac, B. Centromere identity from the DNA point of view. Chromosoma 123, 313–325 (2014).

81. Suzuki, Y., Myers, E. W. & Morishita, S. Rapid and ongoing evolution of repetitive sequence structures in human centromeres. Sci Adv 6, (2020).

82. Arora, U. P., Charlebois, C., Lawal, R. A. & Dumont, B. L. Population and subspecies diversity at mouse centromere satellites. BMC Genomics 22, 279 (2021).

83. Thakur, J., Packiaraj, J. & Henikoff, S. Sequence, chromatin and evolution of satellite DNA. Int. J. Mol. Sci. 22, (2021).

84. Feliciello, I., Picariello, O. & Chinali, G. The first characterisation of the overall variability of repetitive units in a species reveals unexpected features of satellite DNA. Gene 349, 153–164 (2005).

85. Feliciello, I., Picariello, O. & Chinali, G. Intra-specific variability and unusual organization of the repetitive units in a satellite DNA from *Rana dalmatina*: molecular evidence of a new mechanism of DNA repair acting on satellite DNA. Gene 383, 81–92 (2006).

86. Cohen, S., Agmon, N., Sobol, O. & Segal, D. Extrachromosomal circles of satellite repeats and 5S ribosomal DNA in human cells. Mob. DNA 1, 11 (2010).

87. Meštrović, N. et al. Structural and functional liaisons between transposable elements and satellite DNAs. Chromosome Res. 23, 583–596 (2015).

88. Louzada, S. et al. Decoding the role of satellite DNA in genome architecture and plasticity-an evolutionary and clinical affair. Genes 11, (2020).

89. Hsu, T. C. & Arrighi, F. E. Chromosomes of *Peromyscus* (Rodentia, cricetidae). Cytogenet. Genome Res. 7, 417–446 (1968).

90. Greenbaum, I. F. & Baker, R. J. Determination of the primitive karyotype for *Peromyscus*. J. Mammal. 59, 820–834 (1978).

91. Romanenko, S. A., Perelman, P. L., Trifonov, V. A. & Graphodatsky, A. S. Chromosomal evolution in Rodentia. Heredity 108, 4–16 (2012).

92. de Freitas, T. R. O. Ctenomys lami: The highest chromosome variability in *Ctenomys* (Rodentia, Ctenomyidae) due to a centric fusion/fission and pericentric inversion system. Acta Theriol. 52, 171–180 (2007).

93. Baverstock, P. R., Watts, C. H., Hogarth, J. T., Robinson, A. C. & Robinson, J. F. Chromosome evolution in Australian rodents. II. The Rattus group. Chromosoma 61, 227–241 (1977).

94. Lieberman-Aiden, E. et al. Comprehensive mapping of long-range interactions reveals folding principles of the human genome. Science 326, 289–293 (2009).

95. Cheng, H., Concepcion, G. T., Feng, X., Zhang, H. & Li, H. Haplotype-resolved de novo assembly using phased assembly graphs with hifiasm. Nat. Methods 18, 170–175 (2021).

96. Cheng, H. et al. Haplotype-resolved assembly of diploid genomes without parental data. Nat. Biotechnol. 40, 1332–1335 (2022).

97. Li, H. & Durbin, R. Fast and accurate short read alignment with Burrows-Wheeler transform. Bioinformatics 25, 1754–1760 (2009).

98. Li, H. et al. The Sequence Alignment/Map format and SAMtools. Bioinformatics 25, 2078–2079 (2009).

99. Putnam, N. H. et al. Chromosome-scale shotgun assembly using an in vitro method for long-range linkage. Genome Res. 26, 342–350 (2016).

100. Durand, N. C. et al. Juicer provides a one-click system for analyzing loop-resolution hi-c experiments. Cell Syst 3, 95–98 (2016).

101. Li, H. Minimap2: pairwise alignment for nucleotide sequences. Bioinformatics 34, 3094–3100 (2018).

102. Flynn, J. M. et al. RepeatModeler2 for automated genomic discovery of transposable element families. Proc. Natl. Acad. Sci. U. S. A. 117, 9451–9457 (2020).

103. Fu, L., Niu, B., Zhu, Z., Wu, S. & Li, W. CD-HIT: accelerated for clustering the next-generation sequencing data. Bioinformatics 28, 3150–3152 (2012).

104. Hubley, R. et al. The Dfam database of repetitive DNA families. Nucleic Acids Res. 44, D81–9 (2016). 45

105. Smit, A., Hubley, R. & Green, P. RepeatMasker. https://www.repeatmasker.org/.

106. Išerić, H., Alkan, C., Hach, F. & Numanagić, I. Fast characterization of segmental duplication structure in multiple genome assemblies. Algorithms Mol. Biol. 17, 4 (2022).

107. Flusberg, B. A. et al. Direct detection of DNA methylation during single-molecule, real-time sequencing. Nat. Methods 7, 461–465 (2010).

108. Vollger, M. R., Kerpedjiev, P., Phillippy, A. M. & Eichler, E. E. StainedGlass: interactive visualization of massive tandem repeat structures with identity heatmaps. Bioinformatics 38, 2049– 2051 (2022).

109. Jeffares, D. C. et al. Transient structural variations have strong effects on quantitative traits and reproductive isolation in fission yeast. Nat. Commun. 8, 14061 (2017).

110. Liao, W.-W. et al. A draft human pangenome reference. Nature 617, 312–324 (2023).

111. Heger, A., Webber, C., Goodson, M., Ponting, C. P. & Lunter, G. GAT: a simulation framework for testing the association of genomic intervals. Bioinformatics 29, 2046–2048 (2013).

112. Croux, C. & Rousseeuw, P. J. Time-efficient algorithms for two highly robust estimators of scale. In Computational Statistics 411–428 (1992).

